# SubCellSpace: Automated characterization of subcellular mRNA localization patterns in spatial transcriptomics

**DOI:** 10.64898/2026.04.28.720613

**Authors:** David Wouters, Jose Ignacio Alvira Larizgoitia, Nynke Tilkema, Paulien Van Minsel, Ceyhun Alar, Nick Seeuws, Ilias Theodoros Koulalis, Minkyoung Lee, Niels Vandermeulen, Srivathsan Adivarahan, Sandrine Da Cruz, Katy Vandereyken, Andreas Moor, Bernard Thienpont, Thierry Voet, Alejandro Sifrim

## Abstract

The localized translation of transcripts is a universal phenomenon across biological domains. Many examples of subcellular RNA localization and their functional importance have been described. However, these examples remain anecdotal, and a more systematic genome and cell-type-wide analysis is needed. Current spatial transcriptomic techniques can characterize hundreds to thousands of transcript species at subcellular resolutions, enabling the large-scale investigation of subcellular mRNA localization. Here we describe SubCellSpace, a computational framework to learn general representations of mRNA localization patterns. By embedding observed single-cell subcellular localization patterns (SLPs) to an interpretable latent space, SubCellSpace can detect and statistically infer the presence of SLPs, uncover colocalizing gene-pairs and characterize cellular heterogeneity for pattern-presentation. We benchmark SubCellSpace in both synthetic and real data, showing it can correctly detect previously described apical/basal polarized genes in the enterocytes of mouse small-intestine, as well as encode the enterocytes’ orientation. Additionally, we provide a tailored spatial transcriptomics validation dataset for benchmarking SLP identification based on transcripts previously described to be enriched near subcellular structures in HEK293T cells. We propose a practical and computationally-efficient classification workflow that automatically detects localized transcript species and quantifies their degree of patterning, while controlling false positive rates. Finally, we showcase SubCellSpace in both supervised and unsupervised settings, to either classify pre-determined SLPs or to explore spatial patterning without specifying pattern types *a priori*. Automated AI models such as SubCellSpace and their integration in spatial transcriptomics analysis workflows will help characterize previously undiscovered subcellular RNA localization phenomena, providing novel insights into post-transcriptional regulation mechanisms.

## Introduction

RNA localization plays an understudied role in the regulation of gene activity. Transportation of mRNA molecules to specific locations in the cell, mediated by RNA-binding proteins (RBPs) (1), can achieve fine-grained temporal and/or spatial control of protein translation, fast dynamic responses, and thermodynamic energy gains (2). This is particularly important for multi-protein complexes, that need to coordinate expression and assembly of components in time and space (3). Localization is widely observed in prokaryotic as well as eukaryotic biology, playing a critical role in fundamental processes such as cell division and signaling. For example, in Drosophila Melanogaster, *Nanos* mRNA localization to the posterior pole protects it from Smaug-mediated deadenylation, which degrades unlocalized transcripts throughout the bulk cytoplasm, ensuring Nanos protein accumulates only where necessary. (4). In the mammalian gut, the apical-basal polarization of the enterocytes lining the villi is created through specific polarization of both mRNAs and proteins. Local translation of transporter proteins on the apical side of the cell can speed up the intake of nutrients upon feeding (5). In motor neurons RNA molecules are localized to enable rapid cellular responses to external signals, and also play a crucial role in axonal pathfinding and growth-cone steering during development (2). Dysregulation of localization processes can also play a role in disease physiology. For example, mutations in FUS and TDP43, both RNA-binding proteins involved in RNA transport, are the most prevalent genetic causes of amyotrophic lateral sclerosis (ALS) (6,7).

While these studies (2) anecdotally highlight the importance of RNA localization in cellular biology, a systematic study of its constituents and mechanisms, and their role in pathophysiology, remains lacking. Such a study would require high-throughput assays (w.r.t. different genes, cell types and tissues) as well as the computational methods to automate the identification and characterization of such patterns. Current spatial transcriptomics technologies can assay the spatial distribution of mRNAs in high-multiplex at subcellular resolutions (8–10). However, no ST datasets have yet been reported to systematically investigate SLPs by enriching the assayed gene panels with genes previously described and/or predicted to be localized. Due to the scale of current ST datasets, comprising hundreds of thousands of cells across hundreds to thousands of genes, well-performing and computationally efficient methods are necessary to automate SLP discovery. Current state-of-the-art methods have mostly focused on the characterization of FISH stainings at small scales, (11–13), often using manually-engineered data features (i.e. distance to nucleus, nearest neighbor spot distances). This constrains model performance to the inherent limitations of those features (14). Several recent works have evolved to work on more realistically-scaled ST-datasets, but have mostly focussed on gene-colocalization (15,16) or have only been demonstrated in simulations under unrealistic assumptions of very large RNA counts (i.e. minimal n=100) compared to real-life scenarios (i.e. average of n=10) (17). In addition, many of these works take a strictly supervised approach narrowing their scope to a predetermined set of pattern types (e.g. cytoplasmic/nuclear/membrane-bound). Recently, ELLA was proposed (18) which tackles several of these shortcomings and can flag genes as statistically localized using just the nucleus centroid. This method projects cells to a shared, circular coordinate system and statistically infers non-random distributions of transcripts along the nuclear-to-cell-boundary axis. Such mapping is however unlikely to accurately identify certain pattern types (i.e. uniform distribution across a single side of the cell), and testing for a single gene can easily take up to several hours on a standard-sized ST dataset (19–21). More recently, Novoselsky et al. introduced HiVis, a framework tailored to sequencing-based ST platforms (primarily 10X Genomics VisiumHD) (22). HiVis uses image-analysis approaches to assign 2×2 μm bins to predefined subcellular compartments (e.g. apical vs basal, nuclear vs cytoplasmic), and then identifies localized transcripts via differential expression between these compartments. This strategy effectively compensates for the lack of single-molecule resolution in sequencing-based ST and can yield conserved, tissue-level polarization and retention signatures. However, because compartment definitions must be specified *a priori* and quantification is aggregated across cell populations, HiVis is less suited to imaging-based ST data, where individual transcripts are resolved and where cell-to-cell heterogeneity in pattern presentation, unanticipated pattern types, and gene-pair colocalization become tractable questions.

Here, we introduce SubCellSpace, a self-supervised variational convolutional autoencoder that embeds each cell–gene observation from an imaging-based spatial transcriptomics dataset into a latent space capturing its subcellular localization pattern at single-molecule resolution. In contrast to compartment-driven differential-expression approaches, this embedding is learned without pre-specifying pattern types, enabling the identification of transcripts exhibiting non-random localization, the detection of colocalizing sets of transcripts across cells, and the statistical quantification of cell-to-cell heterogeneity in pattern presentation. We extensively benchmark SubCellSpace using both simulated datasets, a novel Xenium ST dataset assaying genes previously described to be subcellularly localized by APEX-seq (23), as well as real-life in-tissue datasets. We provide SubCellSpace as an open-source package containing an end-to-end pipeline which turns standard output of ST techniques into an AnnData object containing the count matrix, single-cell information and subcellular embeddings. This allows for effortless integration of subcellular mRNA pattern information into already existing workflows.

## Results

Throughout the Results, we refer to a single cell–transcript-species combination as an *observation* — the atomic unit on which SubCellSpace operates — and to the learned latent space itself as *SubCellSpace*. The Results are organized in three parts. We first describe the CVAE architecture and characterize SubCellSpace using simulated data, unseen pattern types, and a small MERFISH dataset, to establish that it captures localization pattern structure while remaining robust to confounders such as cell shape and orientation. We then introduce a classification workflow that aggregates per-cell embeddings of a given transcript, statistically infers the presence of a localization pattern, and quantifies its effect size, and we validate this workflow on a novel Xenium dataset of genes with known subcellular compartment assignments from APEX-seq. Finally, we apply SubCellSpace to MERFISH data of the mouse small intestine to demonstrate its utility for biological discovery, recovering apical–basal polarization of enterocytes and, additionally, inferring each cell’s spatial orientation directly from its position in the latent space.

### SubCellSpace learns a general embedding of localization data and separates patterns into separate clusters

#### Learned embeddings are minimally perturbed by confounding variables

To facilitate systematic study of subcellular patterns in spatial transcriptomics, we trained a convolutional variational autoencoder (CVAE) that embeds mRNA localization data in a latent space **(Fig. 1B)**. For each observation, we generate a 100×100 pixel image visualizing its expression pattern. Using images as data type leverages a convolutional layer’s potential for pattern detection (24). As convolutional approaches show degraded performance with single-pixel point clouds, we apply a gaussian blur prior to embedding in a 15-dimensional latent space. These low-dimensional embeddings can then be used to classify patterns in mRNA localization, as well as other downstream tasks such as quantification of pattern heterogeneity, finding co-patterned gene-pairs and detecting unseen patterns in an unsupervised manner. Due to the unavailability of large-scale, annotated ground truth datasets for mRNA localization patterns (SLP), we created a training dataset consisting of simulated patterns (11). We simulated 9 different classes of patterns (including ‘*random*’, i.e. the absence of a pattern) in 317 different cell templates (**Fig. 1A)**. Each simulated observation consisted of a random number of spots, ranging between 10 to 100, of which 90% adheres to its pattern and the remaining 10% being randomly distributed. These empirically determined thresholds were selected to optimize signal detection: lower values resemble random noise, while higher values would saturate the signal, thereby obscuring patterns. To mitigate overtraining on cell identity, we stratified the train-test split for cell-identity, leaving 64 out of 317 cell shapes out of training for validation and augmented each observation through five random rotational transformations. We also left out 3 pattern-types (*nuclear-edge, foci, protrusion)* to investigate what happens to unseen patterns. Due to the nature of the autoencoder, we expect the reconstruction (i.e. the decoding of an embedding) to resemble the original input image. Examples of each unique pattern shows that reconstructions are indeed blurred versions of their respective inputs, implying that the network has learned key features necessary to reconstruct the patterns without overtraining on individual molecule placement (**Fig 2A)**.

**Figure 1:**
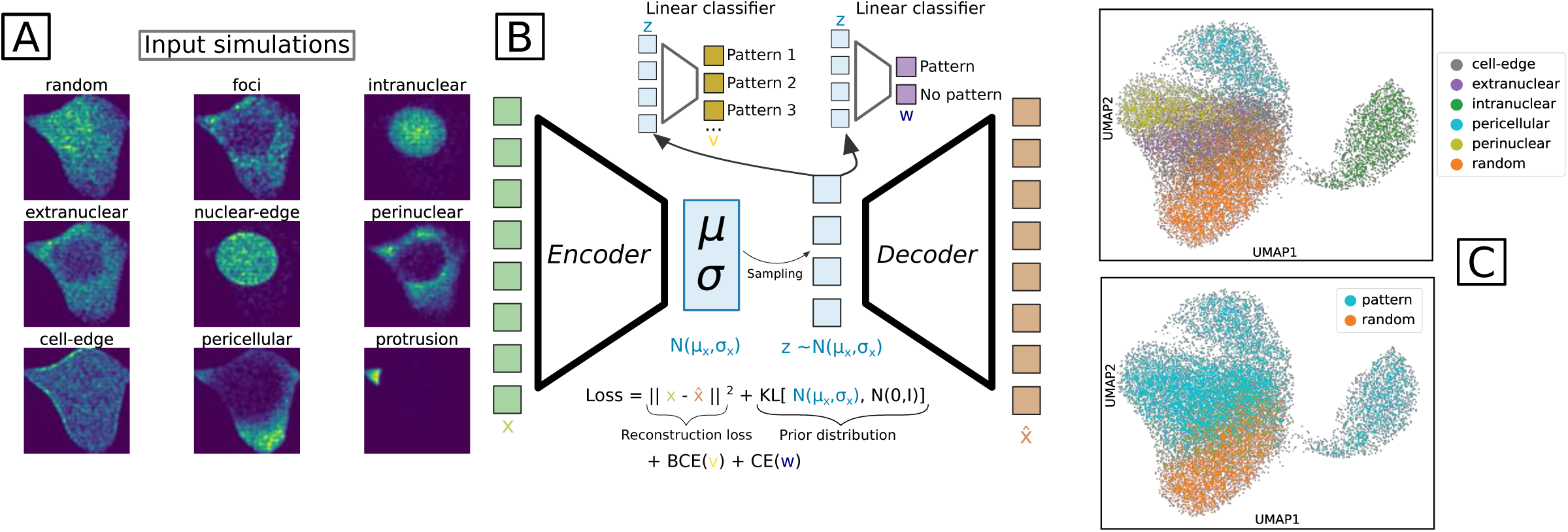
Training of the CVAE. **A.** Examples of each of the 9 patterns available in the simulation framework. Each image shown is a heatmap of 100 simulations of that pattern in a singular cell. **B.** Diagram of the CVAE’s architecture. Blocks represent nodes of the neural network. Trapezoids represent convolutional layers. **C.** UMAP-representation of the SubCellSpace of the simulated validation data, colored by the 2 categories of classifiers contained in the CVAE architecture; unique patterns, and pattern/no-pattern.

**Figure 2.**
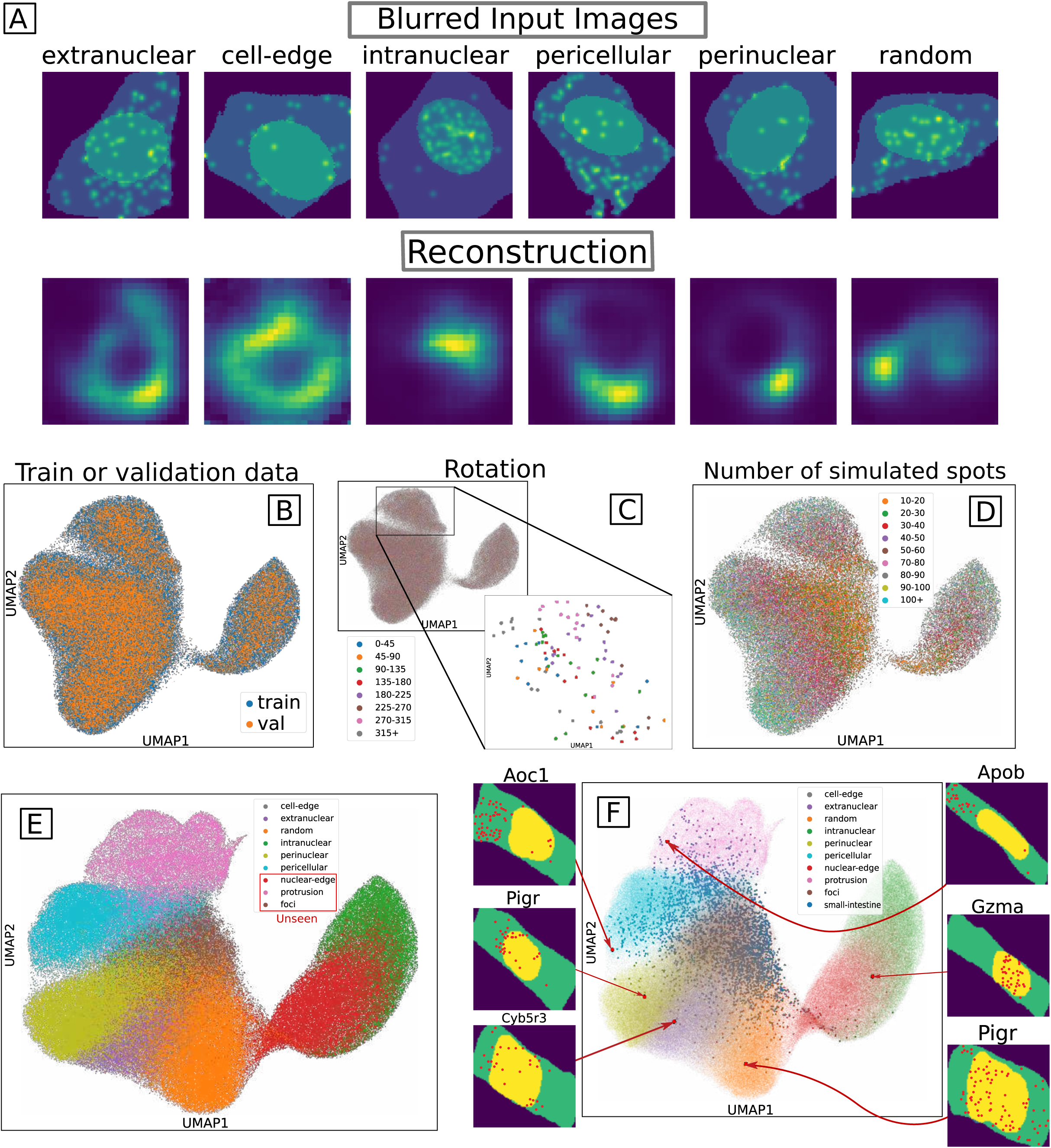
Properties of the learned latent space on seen and unseen data. **A.** Randomly sampled examples of the training data. The top row shows the way observations are fed into the model. The bottom row shows the same data points after a full pass through the CVAE. **B.** UMAP-representation of SubCellSpace’s entire training and validation simulated data colored by their data-suite of origin, showing clear overlap between the two. **C.** UMAP-representation of the SubCellSpace of the simulated validation, colored by their rotated angle. Blown up is the section corresponding to the pericellular pattern, subset for a single cell, showing very local structure based on rotation. **D.** UMAP-representation of the SubCellSpace of the simulated validation, colored by their number of simulated spots. **E.** UMAP-representation of SubCellSpace, now including embeddings of unseen simulated patterns. **F.** UMAP-representation of SubCellSpace, now including embeddings of both unseen simulated patterns and unseen real MERFISH data (labeled small-intestine). Examples of the most outlying real observations per simulated pattern are shown.

To visually evaluate SubCellSpace, we project its 15 dimensions onto a 2D-UMAP space (**Fig. 1C, 2B-F)**. The UMAP of SubCellSpace’s validation data shows clear separation of most simulated patterns (silhouette score = 0.263). Not surprisingly, c*ell-edge* is the least distinguishable pattern subtype (***Table 1***), since it is visually most similar to *random* (**Fig. 1A)**. We also assessed the informativeness of the embedding by looking at the performance in supervised classification of pattern subtypes using a Random Forest classifier on the SubCellSpace, where we achieve an F1 score of 0.736 on validation data (**Supp. Fig. 1**, **Table 2**). In contrast, we did not observe meaningful separation according to training/testing set, cell identity and rotation angle, indicating good generalization to unseen data. We did observe separation according to the number of simulated input spots, reflecting the uncertainty of the embedding given a low signal (i.e. low spot counts tend to cluster with *random* class simulations). (**Fig. 2B-D**, **Table 1-2**). Randomly rotating the nuclear mask of the training data and re-embedding the perturbed input point shows several misplaced intranuclear and extranuclear datapoints. A closer look shows that a datapoint’s embedding can be transposed from intracellular to pericellular depending on the placement of the nuclear mask. This implies that the addition of a segmentation mask aids the network in drawing attention to a certain structure (**Supp. Fig. 2**). In the classification setting, training/validation set and cell shape identity were poorly predicted, with higher predictive performance for number of spots and rotation angle, recapitulating the UMAP plots (**Table 2**). This was to be expected, as the number of spots and their rotation influence correct reconstruction. These could represent features that are desirable to co-embed, as they might contain biological relevance (i.e. cell polarization angle), and are unlikely to perturb the global separation of localization pattern types.

#### SubCellSpace shows clear separation of seen and unseen simulated patterns and real data

Although our simulated data consisted of discrete pattern types, biological use cases might present novel patterns previously unseen in training. We ascertained SubCellSpace’s ability to cope with unseen patterns by only utilising 6 out of the 9 available patterns (*foci, nuclear-edge, protrusion* left out). Projections of these unseen patterns get allocated to an appropriate position in the latent space: the *nuclear-edge* cluster gets positioned as a transitional cluster between pure *intranuclear* and *random* (**Fig. 2E**). *Protrusion*, being a particularly distinct pattern, receives a unique embedding subspace, isolated from the other patterns. Interestingly, at a local level of variability of the embedding, protrusion class embeddings also capture their respective angle (**Supp. fig 3)**. The *foci* pattern subtype is more difficult to distinguish as these patterns represent a very subtle phenomenon (i.e. translational hotspots) and is visually quite similar to *random* after the application of Gaussian blur. These findings indicate that SubCellSpace successfully embeds SLPs into a meaningful latent space that is able to separate different pattern types. It shows resistance to confounders such as cell shape and can project most unseen patterns, likely to be encountered in real data, onto meaningful locations in embedding space.

**Figure 3.**
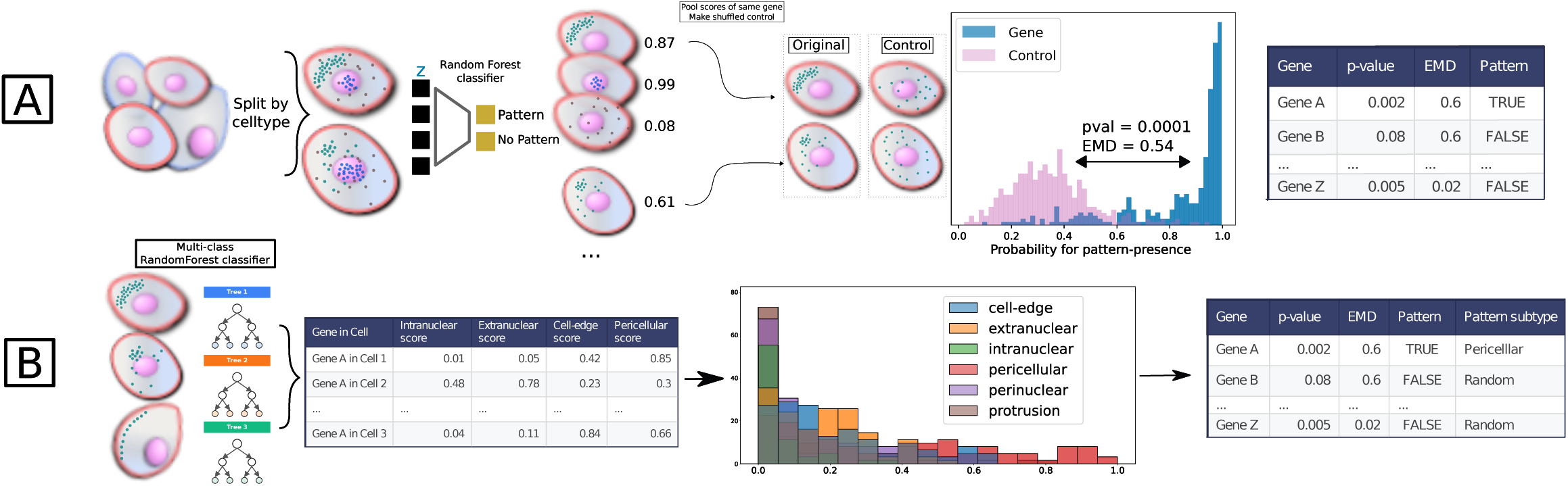
Diagram of the classification workflow. **A.** Diagram visualizing the workflow used to classify an mRNA species as pattern/no-pattern presenting. **B.** Diagram visualizing the workflow to classify positively pattern-presenting mRNA species into pattern-subtypes.

While simulated data offers a way to estimate model performance and structure the embedding space, we simultaneously examined SubCellSpace’s performance on real biological experimental datasets. We applied SubCellSpace to a small (n=408 cells) MERFISH (25) dataset on mouse small-intestine where transcripts are known to be polarized according to the basal-apical axis (5). We specifically focused on the polarized enterocytes and without retraining SubCellSpace, projected patterns within this cell population onto the latent space (**Fig 2F)**. The resulting UMAP shows considerable overlap between the biological data and the simulated patterned data. These real SLPs do not create their own cluster (data points labeled as ‘small-intestine’ in **Fig 2F)**, but instead colocalize with similar simulated data. This also reveals different observations of the same gene projecting to different clusters based on its cell-state at the time of measurement.

Finally, given the drawback of blurring spot signals in the detection of particular pattern subtypes (i.e. *foci*), we explored other neural network architectures specifically tailored to encode *point cloud* data. We implemented a point-cloud autoencoder neural network based on the previously described PointNet encoder (26), and a transformer-like architecture which provides the permutation-invariance inherent to point cloud data. While embeddings of this network are able to accurately reconstruct the input point-clouds and capable of classifying the 6 different patterns based on the simulated data (F1: 0.733, see **Supp. Fig. 4**), separation of patterns (both seen and unseen) within the latent space was much worse (*Seen* silhouette score: 0.0625, *unseen*: -0.078). Additionally, the SI-MERFISH data projected poorly onto the latent space, indicating poor generalizability.

**Figure 4:**
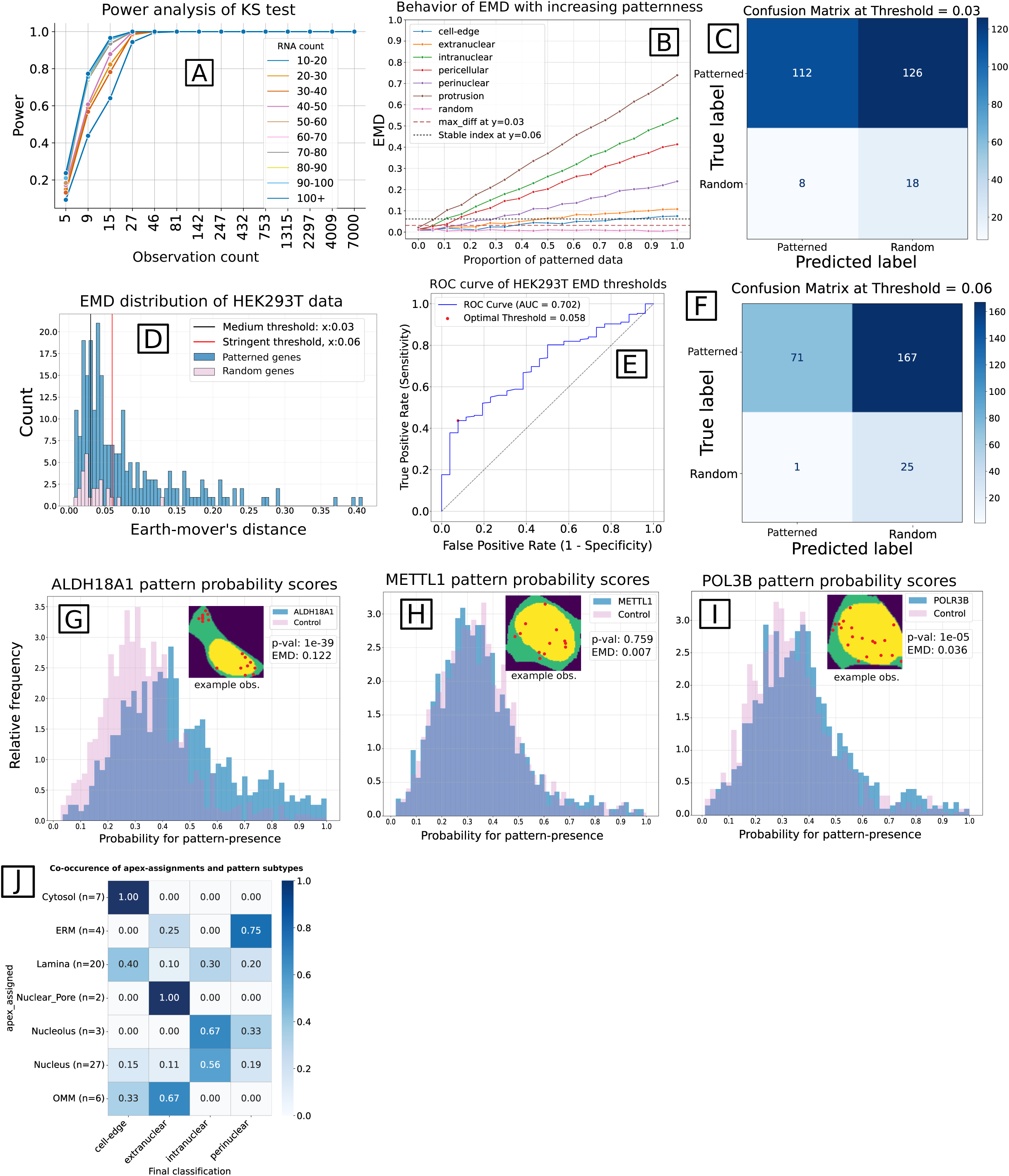
Using SubCellSpace to classify and quantify pattern-presentation in newly generated Xenium validation dataset. **A.** Lineplot showing the results of the power-analysis, split by simulated number of spots. **B.** Scatterplot of the EMD between a mixture (of patterned and random data) and simply random data. The x-axis shows the increasing proportion of patterned data in the mixture, the y-axis shows the corresponding EMD. The scatterplot is split up per simulated pattern. Two horizontal dashed lines represent the two proposed EMD thresholds at y=0.03 and y=0.06. **C.** Confusion matrix showing the results of pattern/no-pattern classification on the APEX-validation Xenium data, using the lenient threshold of 0.03. **D.** Histogram depicting the distribution of calculated EMD’s between all genes from the HEK293T data compared to their randomized counterparts. Histogram split by ground-truth information on pattern/no-pattern presentation. Two vertical lines indicate the two proposed EMD thresholds at x=0.03 (black line) and x=0.06 (red line). **E.** ROC-AUC curve showing the resulting true positive rate over false positive rate depending on the selected threshold. Indicated by a red dot is the optimal threshold, where the area under the curve is highest. **F.** Confusion matrix showing the results of pattern/no-pattern classification on the APEX-validation Xenium data, using the stringent threshold of 0.06. **G-I.** Example of 3 different ways an mRNA species can be characterized by the classification workflow. Each time the classification scores are plotted of the gene’s observations compared to the randomized counterparts, as well as an example observation of the gene. **G**. A clearly patterned gene, with adjusted p-value and EMD that easily passes the set thresholds. **H.** A clearly non-patterned gene, with adjusted p-value and EMD that fall well below the set thresholds. **I.** A dubious gene, whose adjusted p-value barely passes the threshold, but whose EMD indicates low effect size and would not pass the stringent EMD threshold. **J.** Confusion matrix showing the results of pattern-subtype classification on all true positives returned from figure H., split per APEX-assignment. Percentages are normalized across rows.

To summarize, by projecting both unseen simulated and real biological data, the CVAE model demonstrates the ability to effectively embed SLPs. This confirms that the learned latent space meaningfully captures underlying patterns and generalizes beyond the training set to represent the structure of mRNA localization.

### Aggregation of gene embeddings across cells allows the classification and quantification of patterning

#### Probabilistic classification of SLP’s across cells

The findings above describe SubCellSpace’s ability to accurately embed observations of SLPs of individual cells, however ST experiments assay several tens to hundreds of thousands of cells in tissues. In this section, we propose a workflow that aggregates a gene’s embeddings across populations of cells, and both classifies and quantifies patterning. We first train a Random Forest classifier on the simulated data that assigns a probability score to each observation indicating the likelihood of presenting a pattern (i.e. the observation belongs to the *random* class or not). Using this classifier, we score all embeddings of a given transcript, creating a test distribution. To create an empirical null distribution, we randomize each observation by randomly distributing transcript spots across its respective cell-mask. This shuffled dataset is then also embedded and scored, creating the null distribution of random patterning. We use a Kolmogorov–Smirnov test (KS) to determine whether the test and null distributions are significantly different from each other, correcting for multiplicity for each transcript/cell-type combination tested **Fig. 3A**). To estimate the power of this approach, and to estimate the minimal number of required observations to detect several degrees of patterning, we performed a power analysis (**Fig. 4A**). For clearly observable, consistent patterns with low expression number, 8 observations was the bare minimum for detection with a power of 0.5, and therefore chosen as the minimum number of required observations for further analyses. As not every degree of patterning might be biologically meaningful (e.g. very subtle patterning, or patterning present in only a small fraction of cells), we used the simulated pattern-mixture curves to define two candidate Earth-Mover’s distance (EMD) thresholds, corresponding to different stringency levels (**Fig. 4B**). The *lenient* threshold (EMD = 0.03) is placed at the midpoint between the maximum EMD observed for purely random observations and the point at which the weakest pattern subtype (*cell-edge*) curve unambiguously departs from the random baseline, the EMD above which all pattern subtypes are detectable. The *stringent* threshold (EMD = 0.06) is placed at the point where the *cell-edge* curve stabilizes (i.e. no longer decreases as more clearly patterned data is added to the mixture), restricting classification to clear and robust patterns.

#### Classifying subcellular compartment enrichment on novel Xenium validation data

To validate our approach on biological data, we generated a 10X Genomics Xenium dataset based on genes reported to be enriched at distinct subcellular compartments using APEX-seq on HEK-293T cells (23). Using these previously reported significantly localized genes and publicly available bulk-RNA sequencing data, we selected genes that map to each subcellular compartment (i.e. positively patterned genes), as well as expressed genes that map to no specific compartment (i.e. negative controls). Our workflow using the lenient threshold (y=0.03) achieved an F1 score of 0.63, with a precision of 0.93 and a recall of 0.47 (**Fig. 4C**). While we identify many of the reported localized genes, the workflow still reports several false positives. The more stringent threshold results in an F1 score of 0.46, consisting of a precision of 0.99 and a recall of 0.30 (*Table 3)*. Further examination of the EMD distribution across all genes reveals three modes corresponding to clearly non-patterned, clearly patterned, and ambiguous profiles (**Fig. 4D**), with the boundary between ambiguous and clearly patterned lying near EMD ≈ 0.06. This is further reflected in the ROC–AUC curve, which quantifies the trade-off between true and false positive rates across thresholds and shows the more stringent EMD threshold provides the optimal separation **(Fig. 4E**). This trades off a reduced recall of 0.30 with a much higher precision of 0.99 **(Fig 4F**), effectively controlling the false discovery rate in an exploratory setting. We identify examples of patterning of true positive transcripts **(Fig. 4G**) as well as spatial unpatterned true negative transcripts **(Fig. 4H)**. We also find examples of supposedly unpatterned genes whose pattern-probability distribution shows a slight shift and seems to pass initial KS-testing **(Fig. 4I**). However, stringent EMD thresholding correctly keeps this gene from being classified as a patterned gene, in accordance with the findings from APEX-seq, demonstrating the importance of heuristically determining effect size parameters.

Finally, we assigned each mRNA species to specific pattern subtypes. Building on our previous classification, only genes identified as pattern-presenting were considered. Using a multi-class Random Forest classifier trained on simulated data (**Supp. Fig. 5**, F1 on validation data: 0.85), we derived subtype probability scores for all individual observations. Each gene was then assigned to the subtype corresponding to the highest median score, enabling characterization of distinct expression patterns (**Fig. 3B**). A heatmap comparing reported subcellular compartments from APEX-seq (23) with SubCellSpace-assigned pattern subtypes reveals clear concordance: genes localized to the same compartment tend to be classified under the same, or at least a closely related, pattern subtype **(Fig. 4J)**. For example, all cytosol-assigned genes were classified as cell-edge presenting. The ERM-assigned genes were split into extranuclear and perinuclear classifications, reflecting a distinction between diffuse distribution around the nuclear envelope and focused localization at discrete sites along the nuclear periphery respectively (**Supp. Fig. 6**). Genes reported to be nucleus-related were all divided amongst extranuclear, intranuclear and perinuclear, with the core-nuclear assignments (*Nucleus and Nucleolus*) being more classified as intranuclear, and the ones closer to the nuclear membrane being classified more as extranuclear and perinuclear (*Lamina, Nuclear Pore*).

**Figure 5:**
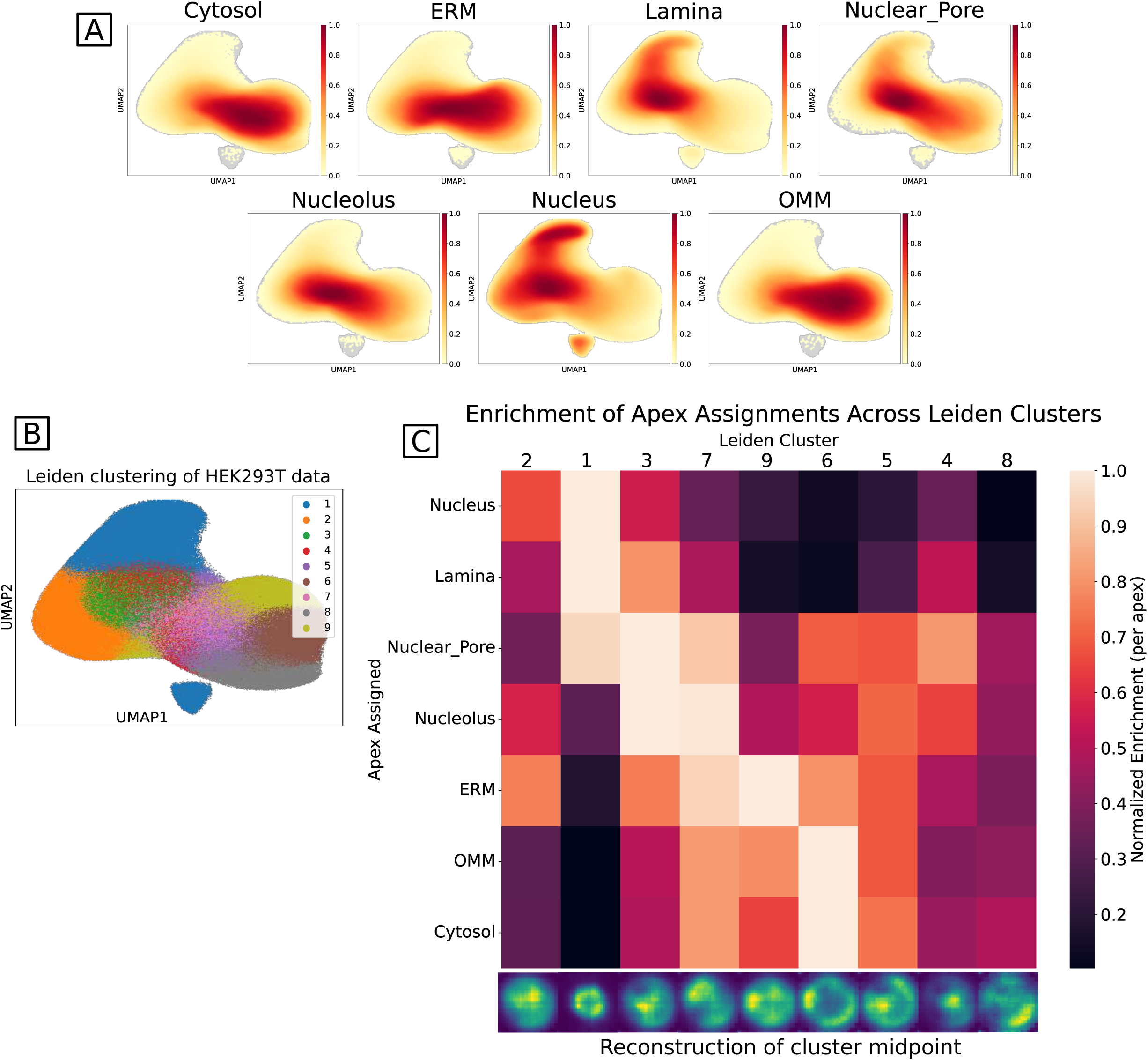
Unsupervised analysis of the APEX-validation data’s SubCellSpace. **A.** A density map of the UMAP-representation of the APEX-SubCellSpace, grouped by each APEX-assignment. **B.** The same UMAP-representation, now colored by a Leiden clustering of resolution 0.6. **C.** Row-normalized heatmap showing the enrichment of APEX-assignments across the Leiden clusters. X-axis labels show the reconstruction of the midpoint of that cluster.

**Figure 6.**
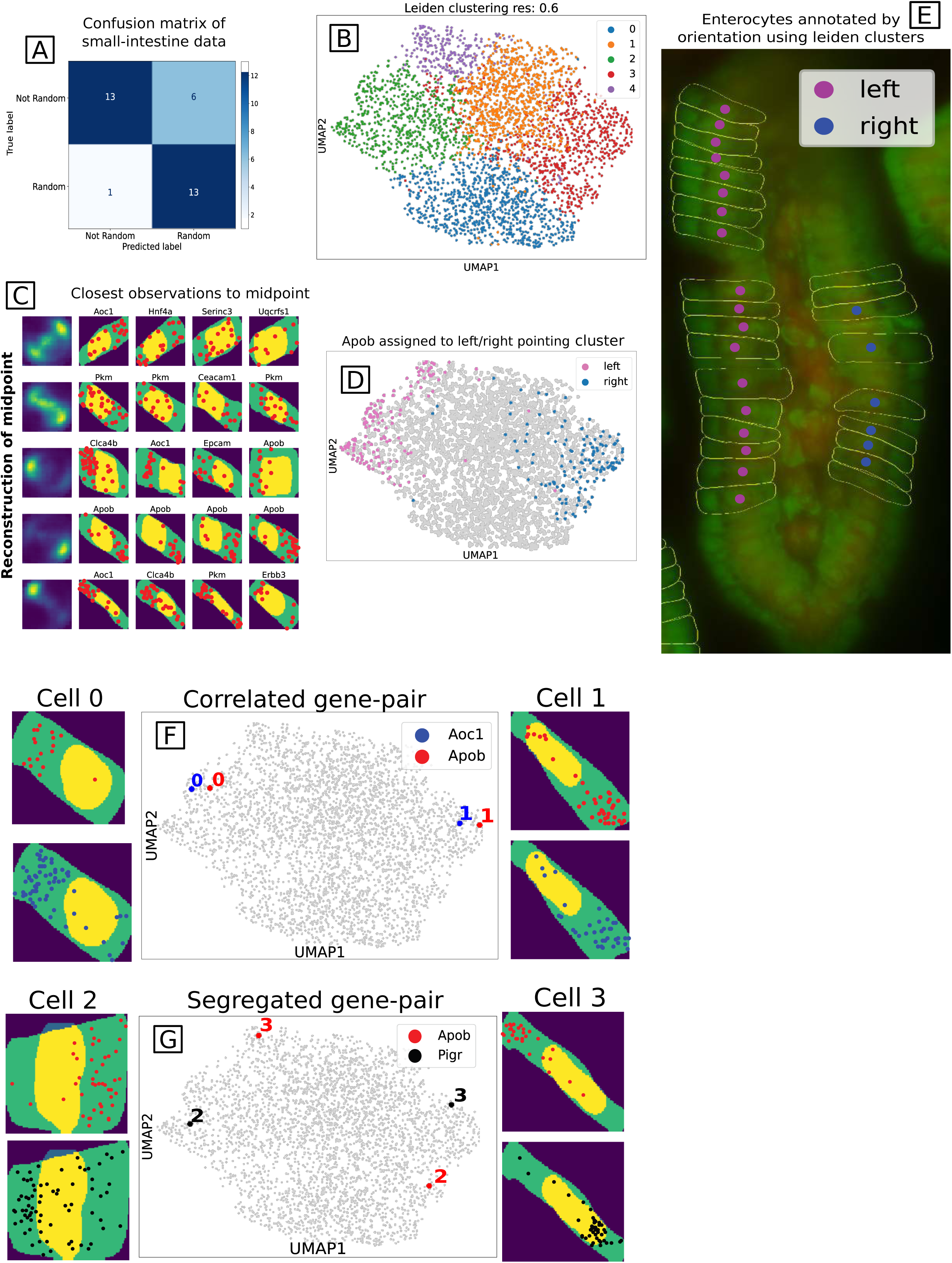
Identification of apical/ basal polarization in mouse enterocytes using small-intestine MERFISH data. **A.** Confusion matrix showing the results of pattern/no-pattern classification on the small-intestine MERFISH data, using the stringent threshold of 0.06. **B.** UMAP representation of the small-intestine’s SubCellSpace, colored by a Leiden clustering with resolution 0.6. **C.** Reconstructions of the midpoint of each Leiden cluster, followed by the 4 closest neighboring observations. **D.** UMAP representation of the small-intestine’s SubCellSpace. Every Apob observation is colored by its presence in either a left-pointing or a right-pointing Leiden cluster. **E.** A subsampled image of the MERFISH stainings. Green staining is the polyT staining, while red is the DAPI staining. Cell segmentation is shown using yellow boundaries, and each DAPI-center is colored by whether that cell’s Apob observation is present in a left-or-right pointing leiden cluster. **F.** An example of the most correlated gene to Apob (Aoc1). Show are 2 cells in which both gene’s expression is drawn. In the middle is the same UMAP as in B and D. Highlighted are the Aoc1 and Apob observations of each cell (indicated by their numbers 0 and 1). **G.** This shows the same figures as panel F, but now with Apob’s most segregated gene: Pigr.

#### Unsupervised investigation of SLP’s

This supervised approach is limited to the composition of its training data, as shown by the lack of direct correspondence between the cellular compartments measured using the APEX-seq technology and the simulated patterns. Because of this, we also explore a more unsupervised approach and interrogate the SubCellSpace directly. While a UMAP-representation shows different localization trends for each APEX-assignment (**Fig. 5A**), they do not perfectly correspond to a Leiden clustering of this space (**Fig. 5B**). These clusters can be interpreted using the generative properties of the variational autoencoder to generate the localization pattern of the cluster centroid. Even though each APEX-assignment shows distinct enrichment profiles across the cluster, there is no clear 1-to-1 mapping between them. Instead, the clusters seem to capture heterogeneity in how an APEX-assignment presents itself, being influenced by cellular factors such as orientation and organelle organisation **(Fig. 5C**).

### Identifying apical/basal polarization in mouse enterocytes

We show the biological relevance of SubCellSpace by revisiting the small-intestine MERFISH dataset in order to assess patterning across the population of observed enterocytes. Our classification workflow correctly identifies 13 of the 19 polarized genes as significantly pattern-presenting. All 6 false negatives pass the EMD threshold, but do not pass the KS-test. While for 2 genes this is likely due to their number of observations (*Cdh1*: n_obs=8, *Ltbr*: n_obs=10) being close to the minimum needed to statistically infer patterning (min = 8, see Power Analysis), the remaining genes’ classification scores indeed do not confidently indicate the presence of patterning (**Supp. Fig. 7**). Conversely, only 1 out of the 14 non-polarized genes assayed were identified as pattern-presenting, yielding an overall F1 score of 0.79 consisting of a precision of 0.93 and a recall of 0.68 (**Fig 6A**, **Table 4**). These results demonstrate that our proposed classification scheme is intentionally conservative, favoring maximal confidence in positive calls over broader inclusion. Additionally, the EMD of the 14 apically-polarized genes show a correlation of 0.77 to the apical bias reported in Lee et al. (27), indicating that the proposed distance metric is a good estimation of their degree of patterning (**Supp. Fig 8**).

However, we can’t immediately designate apical/basal polarization, as this requires cellular information (e.g. which side of the cells point to the lumen). To this end, we first cluster this dataset’s SubCellSpace into 5 clusters **(Fig. 6B)**. Given the distinctive shape of enterocytes, inspection of the midpoints of the clusters as well as their 4 nearest real data neighbors reveals that the first 2 clusters represent dispersed localization signals across the 2 axes. The remaining clusters represent different orientations of polarization along those axes (**Fig. 6C**). Taking *Apob*, a well-known apical-polarized marker, we can annotate cells based on which cluster contains its *Apob* observation. The left/right-pointing behaviour of this SubCellSpace is reflected in its UMAP representation (**Fig. 6D**). Plotting the cellular annotations on the original staining shows that the apical side of the enterocytes have the correct orientation (**Fig. 6E**). Finally, for each cell containing *Apob*, genes located in the same predicted direction as *Apob* (as determined as the cluster assignment) were counted as apical, whereas the opposite direction were counted as basal. Aggregating these observations across cells, we can accurately classify the apical/basal polarized genes by taking a majority vote (**Supp. Fig. 9**). Another approach to identify apical/basal polarized genes is by computing the aggregated cosine distance across cells in the latent space relative to known polarized genes (e.g. *Apob*). In UMAP-space colocalized genes within individual cells will be embedded in close proximity to each other (**Fig. 6F**), whilst genes representing segregated expression patterns will be embedded distantly from each other (**Fig. 6G**). Given this property, we show that even when no known marker gene is available, by manually drawing a pattern, we can embed the resulting images and retrieve genes following that defined pattern. This approach facilitates the exploration of specific spatial patterns without requiring labeled data or a supervised framework (**Supp. fig. 10**).

## Discussion

Spatial transcriptomics platforms enabling the high-throughput characterization of subcellular expression patterns have now become widely available. However, due to the sheer quantity of measured cells and genes, novel computational methods are required to automate the identification of non-randomly spatially distributed transcripts. Current available algorithms often predict simple predefined patterns (e.g. nucleus vs. cytoplasmic vs. membrane-bound) based on manually engineered data features, under strong assumptions (i.e. high expression counts) and have rarely been applied to real experimental data as they often don’t scale to magnitude of modern datasets (i.e. hundreds of thousands of cells over hundreds or thousands of genes). Additionally, no tailored experimentally validated datasets have previously been reported facilitating the development and benchmarking of such methods.

Here we present SubCellSpace, a convolutional variational autoencoder that embeds observations into a latent space representing the diversity of subcellular localization patterns of mRNA species observed. SubCellSpace is able to embed real experimental datasets without the need for retraining or fine-tuning, indicating that the architecture generalizes well to unseen data. The obtained embeddings can be aggregated across cells to probabilistically infer varying degrees of patterning in a cell population within or across samples. To validate the proposed algorithm we generated a novel Xenium Spatial Transcriptomics dataset, using a tailored gene set based on direct proximity labeling by APEX-seq (28). Additionally, we also show the application of SubCellSpace on MERFISH data of small-intestine tissue to automatically characterize polarization patterns.

We leverage the encoded embeddings for supervised classification of patterned versus randomly distributed mRNA species and show good performance in both our validation dataset (Precision: 0.99, Recall: 0.30, F1: 0.46), as well as the detection of known polarized genes in the small intestine (Precision: 0.93, Recall: 0.68, F1: 0.79). By prioritizing confident and robust positive predictions through heuristically determined thresholds, we have effectively controlled for Type-1 error rates in exploratory settings. Beyond simple classification, the framework enables unsupervised exploration, allowing for the detection of co-localized or segregated expression patterns and the characterization of tissue-level features such as apical/basal polarization and cellular orientation. Interestingly, the generative nature of the VAE allows for the identification of patterns even in the absence of known markers, simply by embedding user-defined imagery into the latent space.

Despite good performance on most types of patterns, SubCellSpace is not without technical constraints, most notably in its performance regarding “foci“-type patterns, which consist of randomly distributed clusters of mRNA molecules that are currently obscured by the model’s transformation of discrete points into smoothed signals. However, alternative point cloud-based architectures, which should in theory capture such signals, did not result in improved detections of this type of pattern in our hands (**Supp. Fig. 4**), likely due to the limited number of spots in said patterns. Another limitation of the current architecture is the entanglement of the learned latent space. We observed that several technical factors, such as rotation, spatial orientation and size-related features are embedded across multiple dimensions of the latent space. This behaviour is expected even under β-VAE regularization, which promotes compression and partial disentanglement but does not guarantee alignment between latent axes and semantically meaningful factors. As a result, direct manipulation or correction of specific nuisance variables remains challenging. Future architectures could address this limitation by incorporating stronger inductive biases or structured objectives, such as equivariant architectures that explicitly model rotational or geometric invariances, weakly supervised disentanglement using auxiliary metadata, or contrastive approaches designed to separate invariant biological signals from geometric variability. These strategies may lead to more interpretable latent representations and enable targeted control over specific sources of variation.

The potential applications of SubCellSpace extend beyond mRNA localization into broader multi-omic contexts. For instance, as a proof-of-concept we used transfer-learning on MERFISH experiments with CD45R0 co-staining, with results suggesting the model can effectively detect heterogeneity in protein localization (**Supp. Fig. 11**), however further benchmarking and validation is needed to properly evaluate its ability to cope with subcellular protein abundance patterns. As commercial ST platforms increase the availability of organelle-specific stainings, SubCellSpace could also be extended to focus on structure-specific patterns or translational hotspots by incorporating RBP-specific protein data in an analogous fashion to the one proposed here using the incorporated nuclear DAPI stainings (**Supp. Fig 2**). Applying this framework to cell types with complex morphologies, such as neurons, remains a key objective. Future work focused on adapting the model to cope with irregular cell shapes will be essential for studying mRNA transport in elongated cellular structures, particularly in the context of neurodegenerative diseases like ALS, where distal localization is often disrupted.

In summary, the proposed framework demonstrates how embedding-based representations of spatial transcriptomics data can capture and quantify subcellular SLPs in a unified, interpretable, low-dimensional space. The supervised classification workflow provides a robust means to identify and evaluate known localization signatures, offering a systematic approach for benchmarking and quantitative comparison across datasets. At the same time, the embedding’s structure enables unsupervised exploration of expression behavior, revealing subtle relationships and heterogeneity beyond predefined pattern subtypes. Together, these complementary modes of analysis establish a versatile foundation for both hypothesis-driven and discovery-oriented studies of subcellular organization, bridging the gap between targeted classification and open-ended spatial exploration.

## Materials & Methods

### Dataset preparation

#### Simulated data generation

Data was simulated using the sim-fish package (version 0.2.0) (29). For the training data of the CVAE, a dataset was simulated consisting of the *random, intranuclear, extranuclear, perinuclear, nuclear_edge, cell_edge, pericellular, protrusion* patterns. For each pattern and for each available cell-template (317 unique templates), a single-cell/single-gene observation was simulated by randomly sampling a certain number of spots (ranging between 10 and 100, henceforth termed the *active window*) within the boundaries of the cell, of which 90% adhere to the pattern in question. The coordinates of the spots, as well as the nuclear and the cellular boundary was resized to a 100×100 pixels bounding box; the input size of the CVAE. Input images were then created starting with an empty (meaning all zeros) numpy (version 1.19.5) array of shape (100, 100, 2). Each coordinate pair of the simulated transcriptomic spots were filled in with a 1 in the first (100, 100) slice, which was consecutively blurred using a gaussian blur (*scikit-image.filters.gaussian* (30), version==0.19.3) of sigma=1.5, to enhance its signal. The second slice was used to store the nuclear mask, the cellular mask was kept purely for visualisation purposes but was not included in the training. All input images were then rotationally augmented using a random angle, resulting in a total of 542.425 images. To maintain the (100,100) shape of the input image, we only rotate the content of the image. As this may sometimes cause spots to fall out of bounds, we expand the bounding box where necessary and then reshape it to (100,100).

#### Data processing pipeline

In contrast to simulated data, real ST data requires more preprocessing to get to single-cell/single-gene observations. SubCellSpace features an end-to-end processing pipeline, transforming the standard output of established ST protocols (such as MERFISH or Xenium) into an AnnData object representing its count matrix, enriched with subcellular expression information, allowing for both the visualisation of single-cell/single-gene combinations and their SubCellSpace embedding for downstream analysis.

First, the included images (e.g.: DAPI, membrane staining) are tiled into tiles of 10.000×10.000 pixels, alongside their respective detected transcripts. The transcripts’ coordinates are transposed from physical-units (μm) to image-units (pixels), the method of which depends on the ST-protocol. All following steps are then performed on a per-tile basis. First, cells are (re-) segmented on a combination of the DAPI and the cell-boundary staining using Cellpose’s (version==3.0.10) *cyto2* model with a diameter parameter of 100 (31). The nuclei are segmented separately on just the DAPI using the *nuclei* model. Segmented nuclei are then matched with cell-segmentation, leaving unmatched nuclei out of further analysis. Transcripts are subsequently assigned to the remaining segmented cells. This is turned into a basic AnnData (version==0.10.8) object representing a single-cell-style count matrix. Single-cell information, such as nuclei and cell- segmentation masks, are added to the AnnData object alongside a dataframe containing the detected transcript of each cell. This single-cell information is then used to create input for the CVAE as described above, and embeddings of the CVAE are then added as an extra matrix layer to the AnnData object, enriching the count matrix with subcellular information. Based on the size of the slide and the number of transcripts detected, processing can range from 1.5 hours to 8 hours, with the main bottleneck being the cell-segmentation which is performed on GPU, where parallelisation is constrained beyond its intrinsic capabilities. Analyses can be considerably faster if the default segmentation results are acceptable, thereby eliminating the need for additional segmentation steps.

### Mouse small-intestine enterocytes validation MERFISH data

Raw MERFISH data was obtained from previously published (27). As this data was generated using an older MERFISH protocol, the cell-boundary staining was of too low quality to perform robust large-scale cell-segmentation on using Cellpose. However, due to the distinct shape of small-intestine enterocytes, manual segmentation was easily performed, delineating 408 cells. The rest of this data was processed as described in *Data Processing Pipeline*.

### Xenium validation experiment on HEK293T cells

#### Gene selection

Positively localized genes were selected based on the adjusted logFC change reported by Fazal et al 2019 (23). Per cellular compartment analyzed (8 in total), the top 20 most significantly enriched genes were selected alongside 50 other significantly localized genes, after first verifying a decent expression level in both bulk RNA data (GEO: GSE146991) and single cell data (GEO: GSE197769). From the same datasets, 30 genes of medium expression level were selected that were not significantly localized in HEK293T cells, which serve as control non-patterned genes.

#### Culturing of adherent cells on Xenium slide

HEK 293T cells were cultured in IMDM medium (Thermo Fisher, 12440053) with 10% FBS (Thermo Fisher, A5256701) and 1% penicillin-streptomycin (Thermo Fisher, 15140122) in an incubator at 37°C with 5% CO2. Xenium slides and corresponding cassette (10x Genomics, PN-1000460) were sterilized by submerging in 70% EtOH followed by 30 min UV treatment. Slides were coated with 5 ug/cm2 fibronectin (Sigma Aldrich, F0895) in DPBS (Thermo Fisher, 14190094) for 15 min at 37°C, and left to air dry for 2 hours at room temperature. The coated surfaces were carefully washed three times with DPBS and once with complete media. Cells were harvested using Trypsine (Thermo Fisher, 25200056) and suspensions were counted using the LUNA-FL Dual Fluorescence Cell Counter. A total of 5 × 105 cells were plated per slide and grown overnight.

#### Xenium Assay

Slides were prepared according to demonstrated protocol for Xenium In Situ for Fresh Frozen Tissues – Fixation & Permeabilization (CG000581, Rev D, 10x Genomics) with adaptations for handling of adherent cell lines. Briefly, the slides were washed three times with 750 µl of 1x DPBS and fixed with 4% paraformaldehyde (Electron Miscroscopy Sciences, 15700) for 30 min at room temperature. Permeabilization was performed using pre-chilled 100% methanol (VWR, 20846) at -20°C for 1 hour. After a final DPBS washing step to remove all remaining methanol, the slides were hydrated with DPBS. All remaining steps were performed on the prepared cell culture slides according to the Xenium In Situ Gene Expression – with Cell Segmentation Staining (CG000749, Rev A, 10x Genomics) apart from the autofluorescence quenching step which was omitted. Finally, the cassettes with the fixed cells were loaded on the Xenium analyzer instrument.

#### Processing into AnnData object

The resulting raw Xenium data was processed as described in *Data Processing Pipeline*. Segmentation was thus re-performed using only the 16sRNA staining, as this is often the highest quality one, resulting in 154296 cells that could be matched to unique nuclei.

### Convolutional Variational Autoencoder (CVAE)

#### Architecture

Convolution layers are capable of detecting patterns in image processing tasks. Relying on this, the model utilises 4 consecutive convolutional layers including a ReLu activation and maxpooling step after each layer, followed by a fully connected layer before encoding the final tensor into a 15-dimensional latent space, represented by a 500×15 **μ** and **σ** tensor of equal size. The decoder then followed a similar, but asymmetric deconvolution scheme, resulting in a reconstruction of the input image. The loss was an addition of mean squared reconstruction error and Kullback–Leibler (KL) divergence, which makes this autoencoder variational. This aspect improves generalizability through regularization, and interpretability by allowing the efficient sampling and visualization of the latent space. The reconstruction loss was only calculated on the first slice of the reconstruction, which represents the gene localization. Reconstructing the second slice, representing the nuclear shape, would incentivise the model to learn the cell shape resulting in worse generalisation. In parallel to this architecture, 2 classifiers were implemented. One to classify the embeddings as being pattern-presenting or not, and one to classify the embeddings as presenting a specific pattern. These classifiers used (binary) cross entropy as their loss, where the weight of the binary loss was multiplied by 100, as correctly classifying pattern-presence is more important than correctly identifying which pattern. These losses were both added to the total loss for model training as used for self-updating. All architectures were implemented using PyTorch (version==1.12.1).

#### Training the model

To train the CVAE, 3 patterns (*nuclear-edge, foci, protrusion)* were left out from the simulated dataset described above to be able to investigate the introduction of unseen patterns to the latent space. In addition, a train/validation split was done by stratifying the cell-templates the simulations were based on, leaving 64 cell-templates out entirely, to assess whether the model was overtraining on cell shape rather than localization patterns. The model was trained for 100 epochs, with a learning rate of 1e-05, a separate learning rate of 1e-03 for the classifiers, a beta of 1e-04 as weight for the KL-divergence, and 15 latent dimensions. Both more epochs and more latent dimensions showed an unstable validation loss, indicating that while reconstruction improved on the training data, the model’s generalisability would decrease. Training was performed on an NVIDIA A40, and took approximately 3 hours.

#### Point-cloud based application

An alternative approach to creating the latent space was explored by representing the input data as a point-cloud. The autoencoder is then adjusted according to state-of-the-art point-cloud applications. The encoder converts point clouds of variable size into a fixed-size “feature vector” of size 128. We use a variation of the basic PointNet architecture with 2 T-Net modules in the first, point-specific, stage of the model. As input the model expects a set of points with x-y coordinates and a label indicating whether the point is part of the cell nucleus. A transformer-like architecture decodes the encoded representation back into a point cloud. The transformer structure provides the permutation-invariance inherent to point cloud data. The decoder produces a variable-size output based on a finite set of “reconstruction tokens”, similar to the token embeddings used in language modeling. During training the decoder is provided with the same number of tokens as there are points in the input point cloud. The decoding process produces a reconstruction by combining the tokens with the point cloud representation provided by the encoder. This scheme avoids a need for the encoder to extract information related to the number of points and allows for more focus on pattern-specific details. Because of the transformer decoder we cannot rely on consistent ordering of output points. Losses should reflect this by being permutation-invariant. Permutation-invariance is provided by a “preprocessing” step when computing the losses. The best mapping between input and reconstruction point cloud (best in terms of sum of Euclidean distance between the points) is computed. Total loss consists of the reconstruction loss which computes a point-wise distance between matched points, cross-entropy between binary nucleus labels of the input and classification scores (predicting “nucleus membership”), and a KL-divergence for variational regularization.

### Localization pattern classification and quantification

#### Training a classifier

Unless specified otherwise, all references to training a classifier were performed using the following scheme: available data was divided into train/test data using an 80/20 split. A Random Forest Classifier was trained using bootstrapping of 100 estimators. Parameter tuning of the depth and features used was performed with crossvalidation (sklearn.model_selection.GridSearchCV), using AUC as performance metric. The best performing parameter configuration was then used to train a final classifier, which was then evaluated on the original test data exactly once.

#### Pattern classification and quantification

To create a classifier that delineates patterned from non-patterned data, training data was generated by taking highly patterned simulated data (similar to the training data of the CVAE). Non-patterned data was generated by shuffling the expression of the patterned data within the bounds of the cell and resulting ‘shuffled’ gene expression images were embedded into the SubCellSpace. Both datasets were joined and used to train a Random Forest Classifier which assigns a pattern-probability score to individual gene-cell observations.

Pattern-probability scores for each observation of a gene are then aggregated across cells, and compared to the same scores from their shuffled counterparts. The two probability density functions were then compared using a two-sample Kolmogorov-Smirnov (KS) test (scipy.stats.ks_2samp, version==1.10), evaluating whether they originate from the same underlying (unknown) continuous probability distribution. As an estimator for the effect size of this test, we also calculate the earth mover’s distance (EMD) between distributions, in the form of scipy’s implementation of the wasserstein distance.

The **pattern-subtype** classifier was trained in a similar manner. However, the training data now consists of the *intranuclear, extranuclear, perinuclear, cell_edge, pericellular, protrusion* and *random* patterns. This simulated data was only sampled from the cell-templates not included in CVAE training data. As this reduces the dataset-size, the train-test split was stratified for the *n_spots_interval* variable to ensure equal distribution. While training, each pattern-subtype is treated as its own label, creating a multi-label RandomForest classifier that assigns a probability score for each pattern-subtype to single observations. Finally, a gene is then classified as its highest median pattern-subtype score. This max calculation does not include *random*, as the previous classification round has already determined a pattern is present. It was still included in the training in the assignment of probability scores, since single observations can still be *random*.

#### Power analysis

The power analysis was conducted following the methodology of Baumgartner & John Kalassa (32). Specifically, cell counts (sample sizes) were evaluated for 14 rounded counts on a log10 scale ranging from 5 to 7000. For RNA counts, the binned RNA counts [0-10, 10-20, 20-30, 30-40, 40-50, 50-60, 60-70, 70-80, 80-90, 90-100] were used. For each combination of cell count and RNA count, 1000 random test and control genes were obtained using the simulated gene function and its shuffled counterparts, without a random seed ensuring each gene was unique. For each of the 1000 test-control pairs, test statistics were determined with the two-sample KS test. These results were compared against critical values of 0.05 and 0.00001 (Bonferroni correction for 5000 tests). The number of times the null hypothesis was rejected out of the 1000 samples was counted and then divided by 1000 to determine the power for each combination.

### Latent space analysis

#### Colocalization calculation

Gene-pairs were filtered based on whether there are at least 4 cells in which they are both expressed. Then, for each cell, the pairwise similarity is calculated between the embeddings of gene-pairs present in the cell using Python’s Optimal Transport’s implementation of cosine distance. A gene-pair is then assigned their median distance across all cells.

#### Unsupervised clustering analysis

Each observation was assigned its 25 nearest neighbors using scanpy<u>.pp</u>.neighbors (version==1.9.1). The community Leiden algorithm was then applied on the resulting neighborhood graph with a default resolution of 0.6 unless specified otherwise. The same neighborhood graph was also used alongside the latent space’s top 10 principal components (<u>sc.tl</u>.pca) to calculate a 2D-UMAP representation of the embeddings, to facilitate visualization of SubCellSpace. Visualizing the contents of clusters was done by calculating the 15-dimensional midpoint of all observations in the cluster. This midpoint is treated as a new embedding and decoded as described in *training the model*.

## Data availability

All processed datasets used are available on Zenodo: 10.5281/zenodo.19708995

For the raw small-intestine MERFISH data used, please refer to their original publication (27).

## Code availability

All code necessary to replicate the training of SubCellSpace and its analyses, as well as make the figures shown and apply it to new datasets is available on GitHub.

## Funding

This research was funded by the Leuven Future Fund (LISCO-BIOMED), Opening the Future Fund, KU Leuven ID-N (3E210655 COLUMBO) and Fonds Wetenschappelijk Onderzoek (FWO) DeepSubCell project (G005923N). In addition, co-authors had the following funding support: JIAL: FWO SB: 1SF1522N and 1SF1524N, PVM: FWO 11K6222N, ML: European Research Council (Grant ID: 949717) and the Swiss National Science Foundation (Grant IDs: 190799, 190709), SA and AM: Novartis FreeNovation Grant and the ETHZurich Postdoctoral Fellowship, SDC: FWO G064721N, the Muscular Dystrophy Association (MDA, 962700), KU Leuven Opening the Future and VIB-Tech Watch

## Abbreviations

ST: Spatial transcriptomics
SLP: Subcellular localization pattern
CVAE: Convolutional Variational AutoEncoder
RBP: RNA-binding proteins
KS: Kolmogorov–Smirnov
EMD: Earth Mover’s Distance
SI: small-intestine

---

**Supp. Fig. 1:**
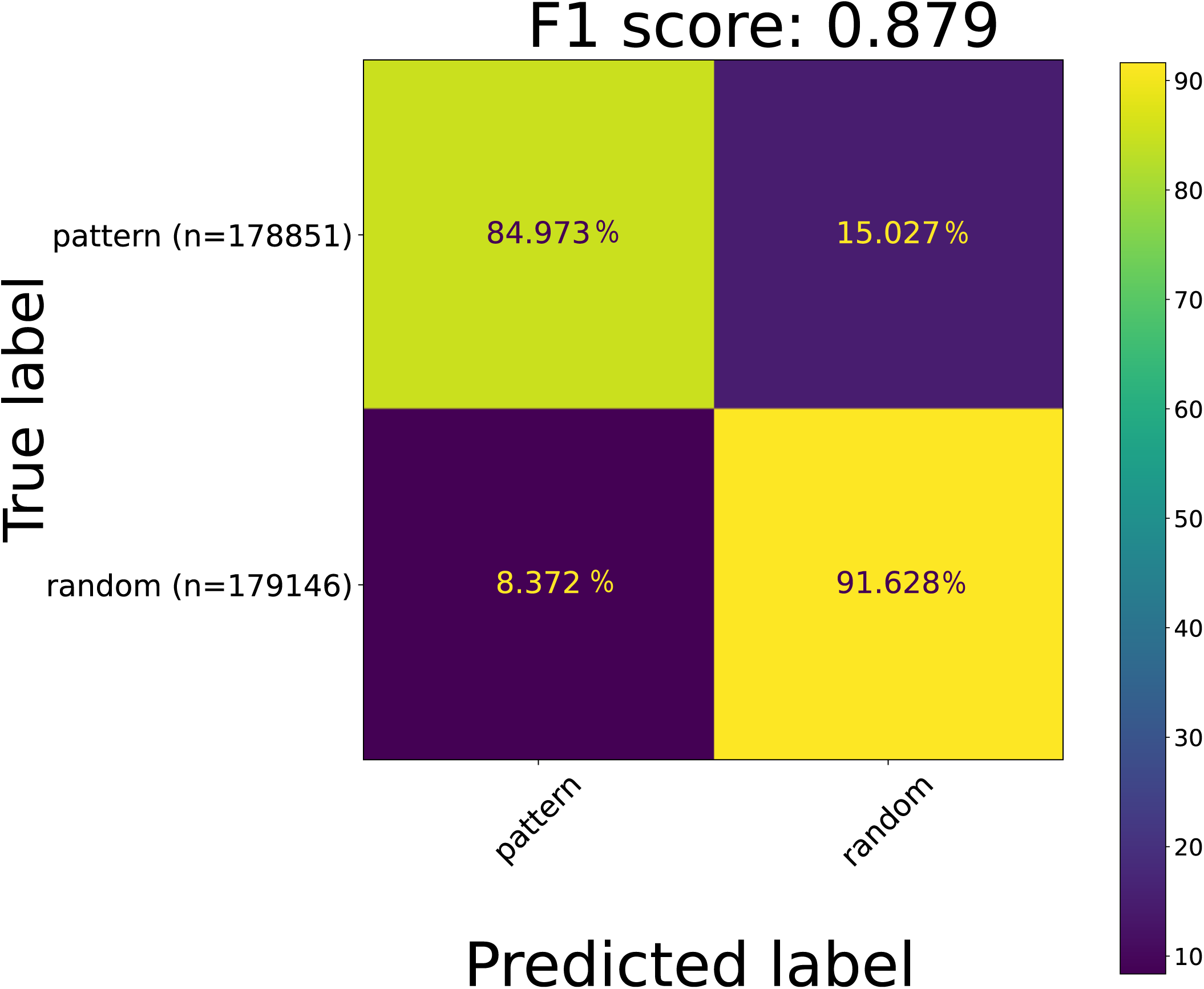
Pattern/no-pattern classification performance on simulated validation data. Confusion matrix of the pattern/no-pattern classifier on the simulated validation data. Cell elements are row-wise normalized percentages. Total *n* is noted next to the true labels.

**Supp. Fig. 2:**
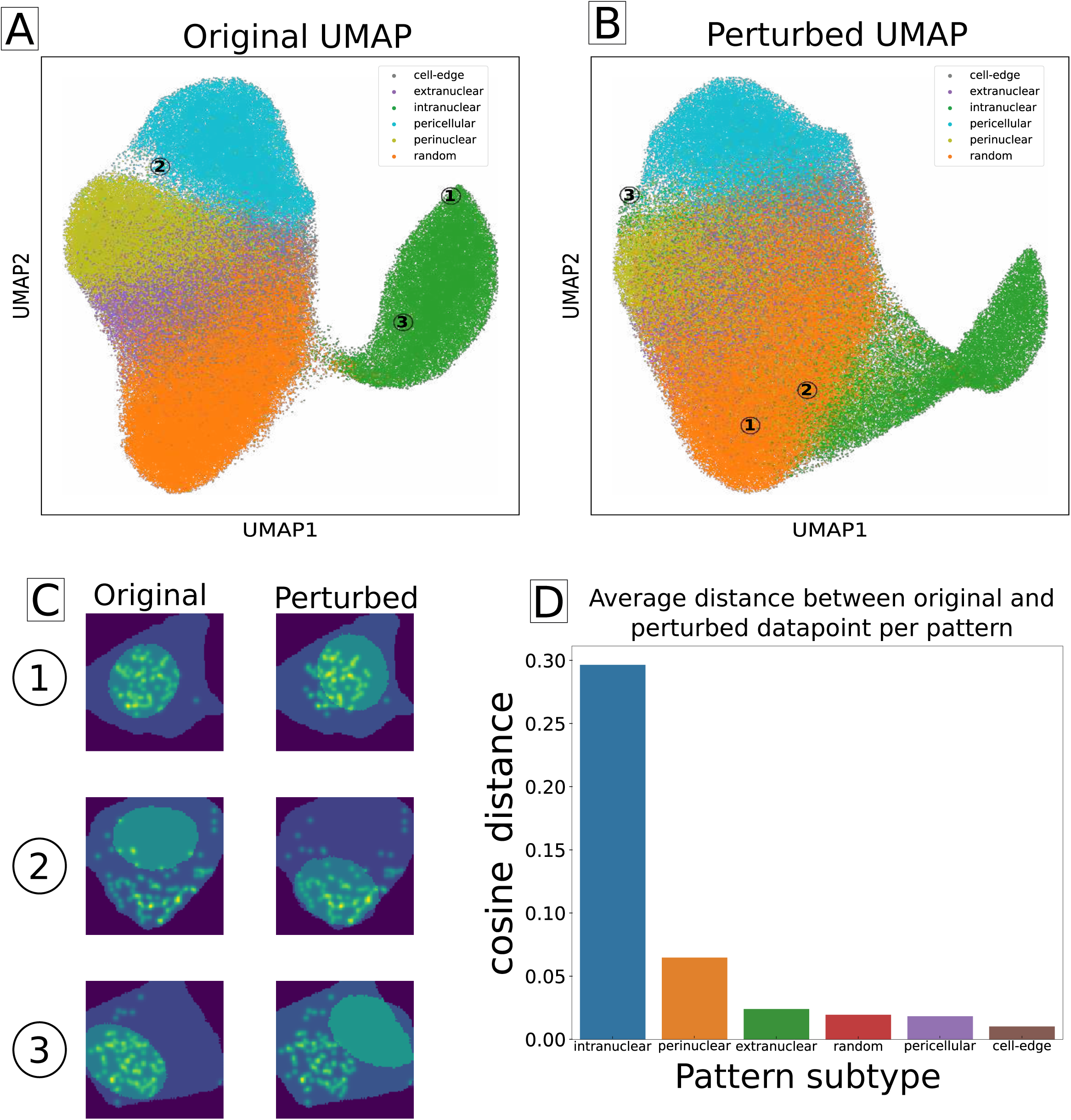
Perturbing the DAPI mask disrupts embeddings of nuclear-based patterns. **A.** UMAP-representation of regular SubCellSpace, with 3 maximal perturbed observations indicated. **B.** UMAP-representation of a SubCellSpace after each DAPI-mask was randomly rotated. The same 3 observations are indicated with their new placement. **C.** 3 maximally perturbed observations, measured by pairwise cosine distance in multi-dimensional space. The left column is their original form, right is after perturbing the DAPI mask. **D**. Averaged distance per simulated pattern between original embeddings and perturbed embeddings.

**Supp. Fig. 3:**
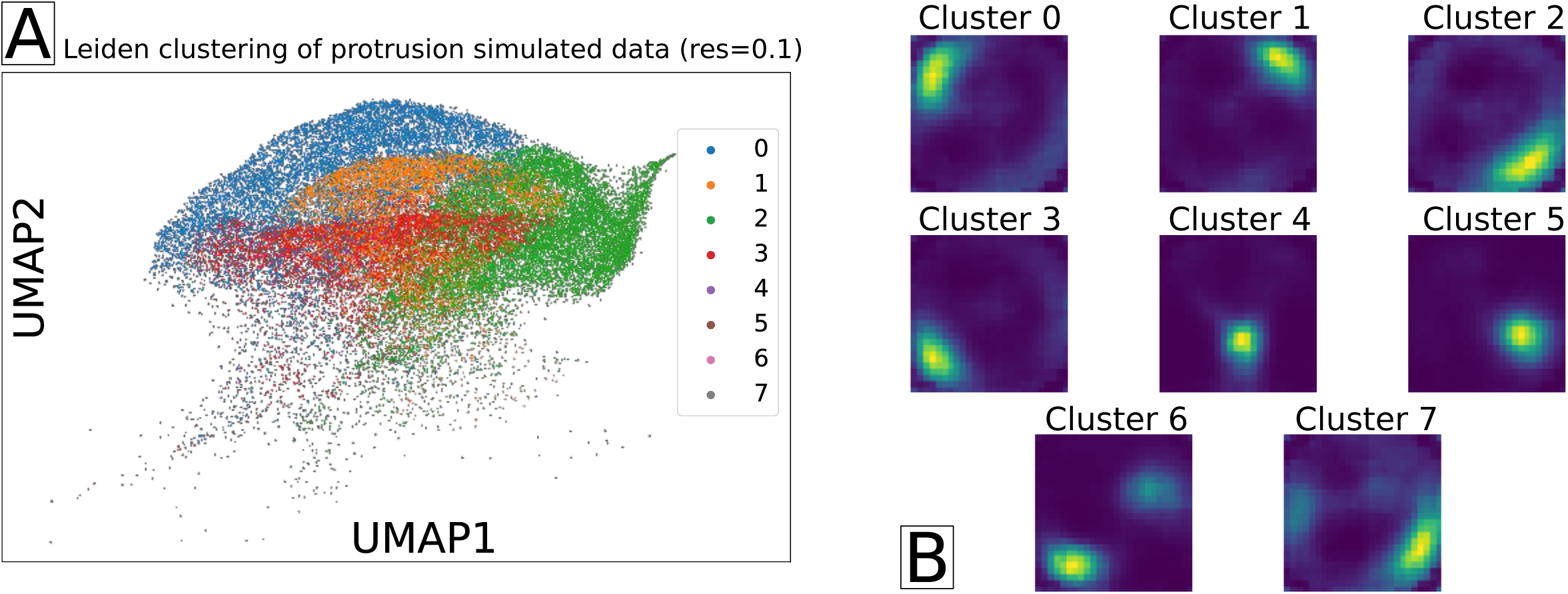
Subclustering of the unseen protrusion simulated pattern. **A.** Cropped UMAP-representation of SubCellSpace showing only the unseen protrusion simulated observations, colored by cluster assignment by a Leiden clustering at resolution 0.1. **B.** Reconstruction of the midpoint of each leiden cluster.

**Supp. Fig. 4:**
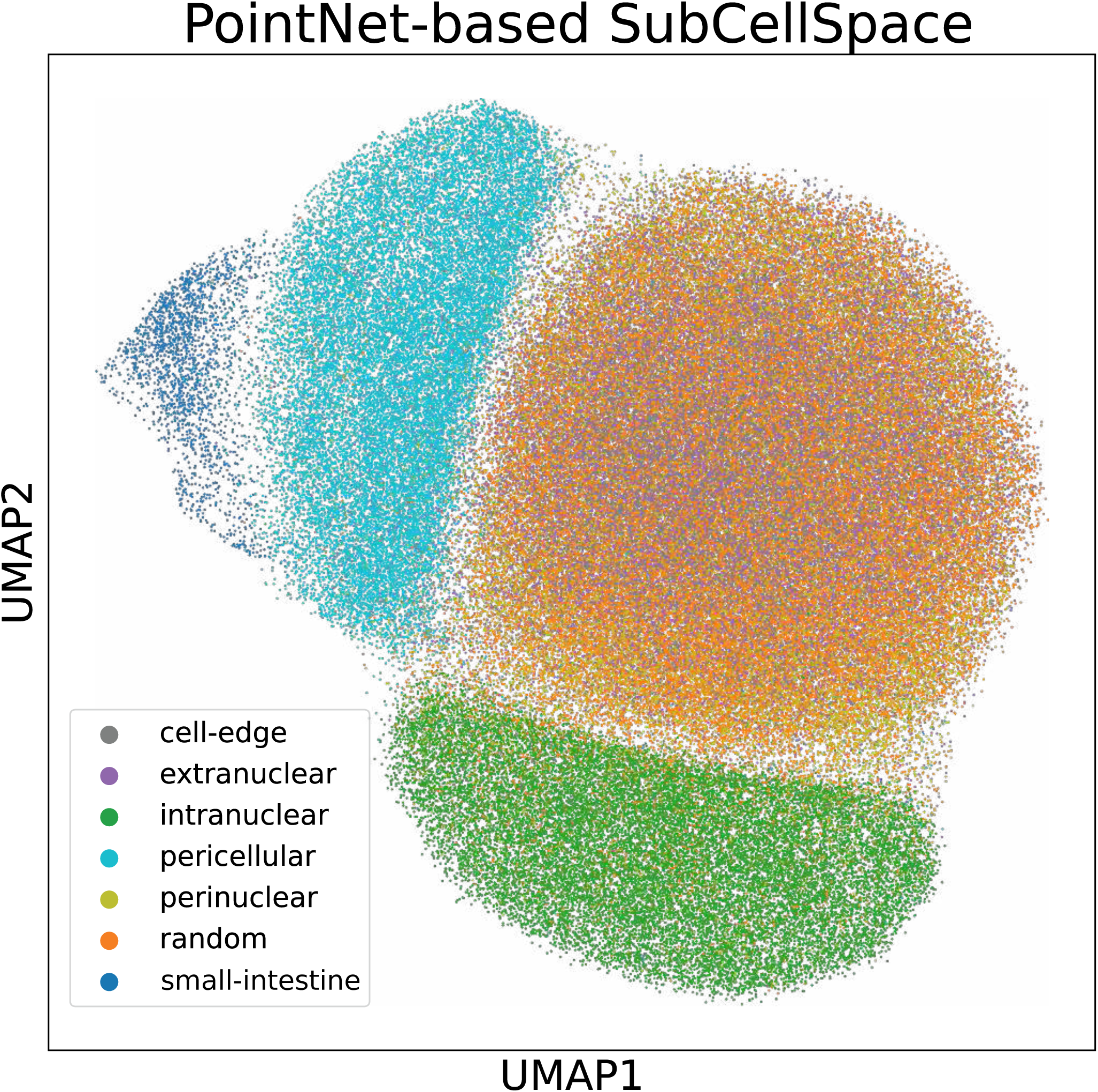
Point-cloud implementation of SubCellSpace. UMAP-representation of the training data combined with the small-intestine MERFISH data. UMAP was calculated based on the point-cloud application-derived embeddings.

**Supp. Fig. 5:**
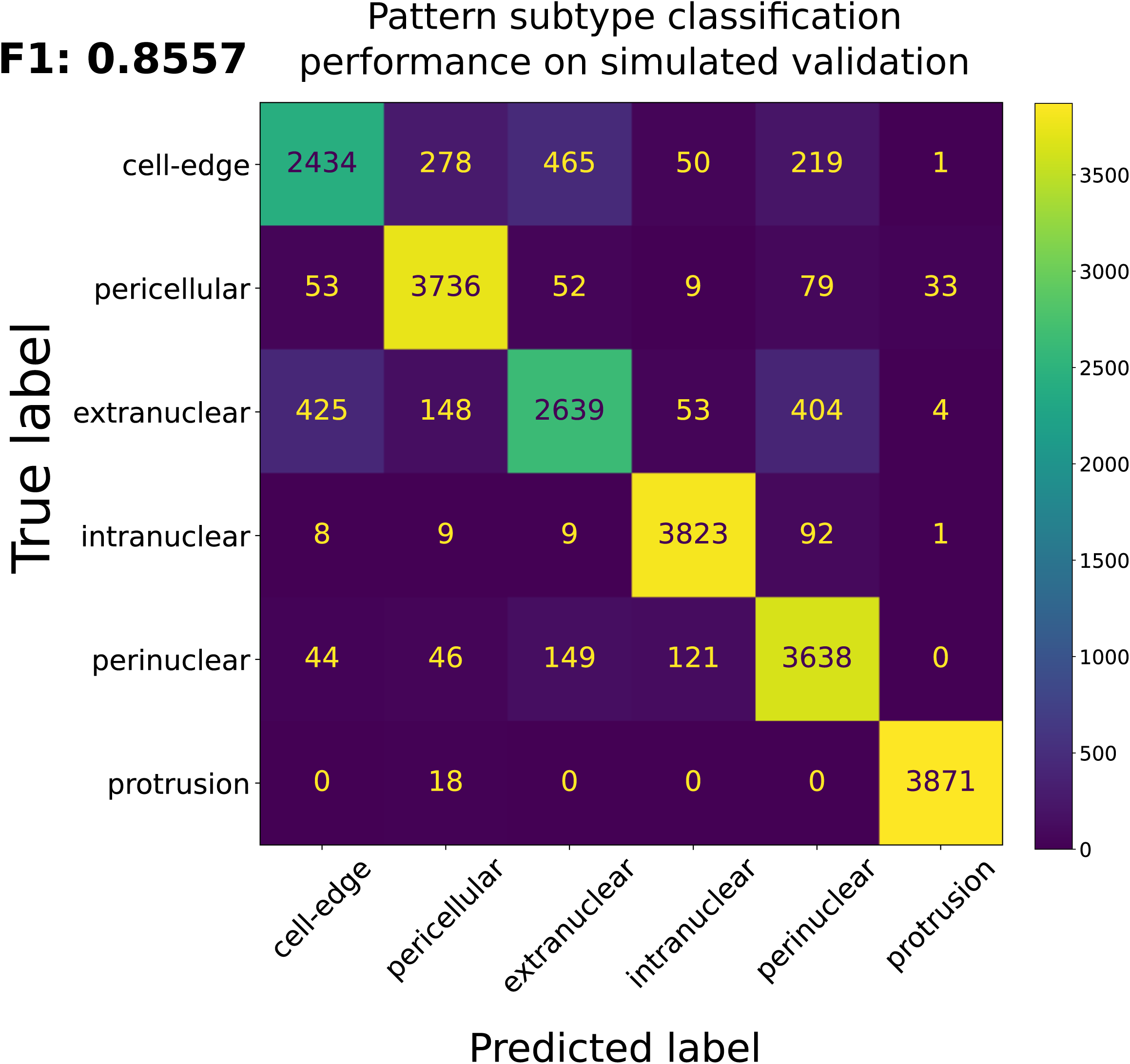
Pattern subtype classification performance on simulated validation data. Confusion matrix of the pattern subtype classifier on the simulated validation data. Cell elements are absolute values.

**Supp. Fig. 6:**
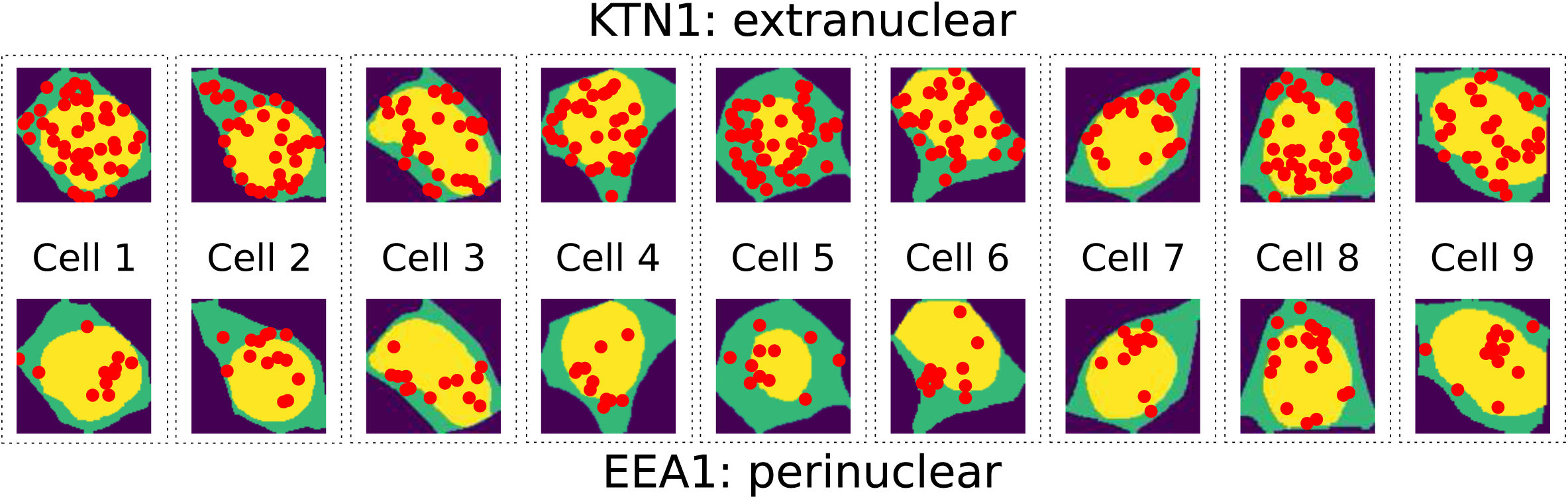
Heterogeneity in pattern-presentation of ERM-assigned genes. Shown are 2 genes from the Xenium validation data, both of which have been assigned to be ERM-enriched by APEX-seq, but classified as different pattern subtypes by our workflow (KTN1 as extranuclear, and EEA1 as perinuclear). Columns show nine cells with maximal respective pattern-subtype scores and their mRNA localization in those cells.

**Supp. Fig. 7:**
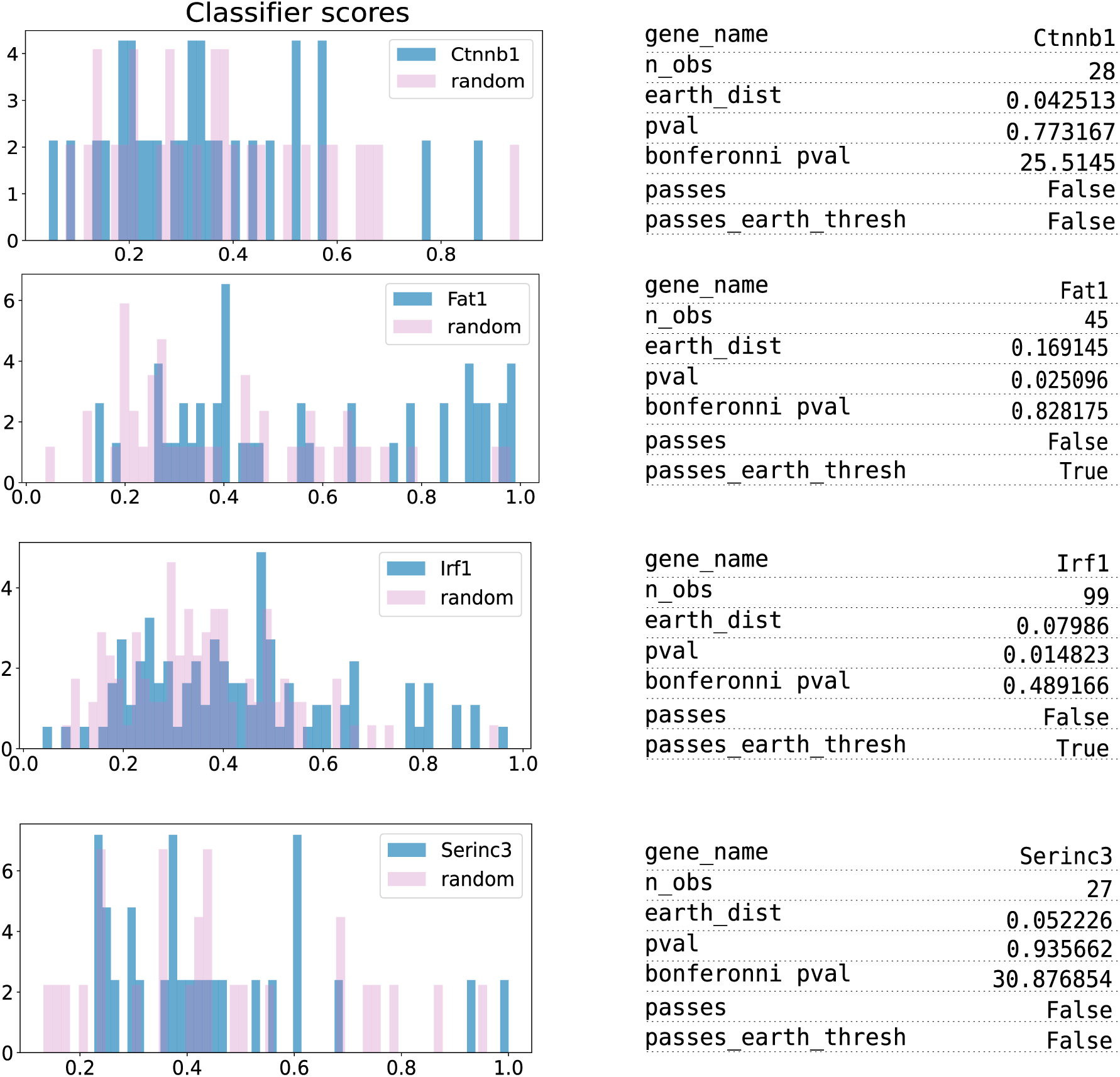
Pattern-probability distributions of False Negatives of the small-intestine MERFISH data. Pattern-probability distributions for each gene of the small-intestine dataset that was falsely classified as not-pattern-presenting, together with its randomized distribution. Noted for each gene is their EMD, p-val, adjusted p-val, and whether those values pass their respective thresholds.

**Supp. Fig. 8:**
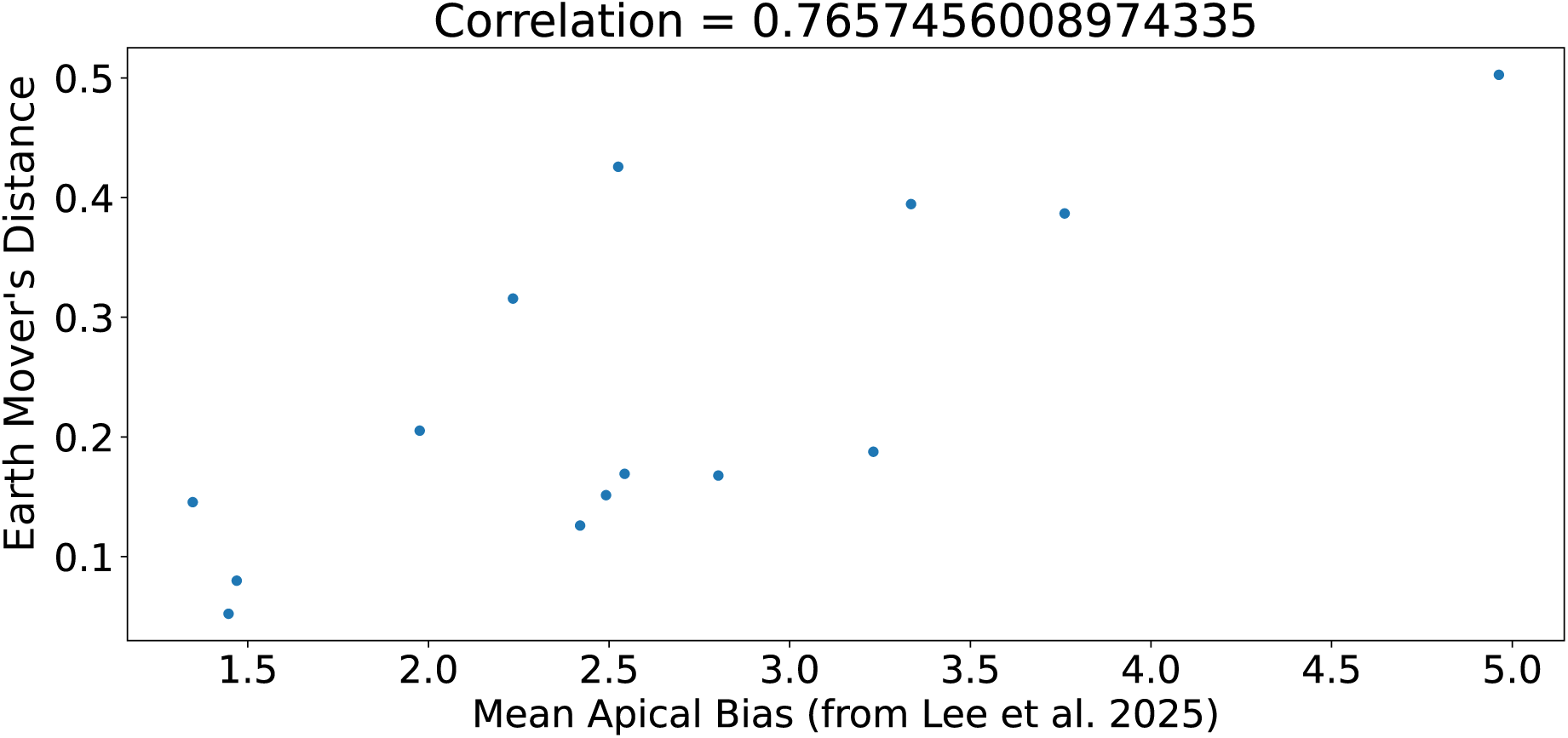
EMD’s of apical genes correlate to their apical bias. A scatterplot showing the correlation for each ground-truth apically polarized gene from the SI MERFISH data. Each dot is a gene, where the x-axis represents its noted apical bias as reported in Lee et al. 2025. The y-axis is this gene’s EMD compared to its randomized counterpart. Correlation was calculated as the Pearson’s correlation between the two metrics.

**Supp. Fig. 9:**
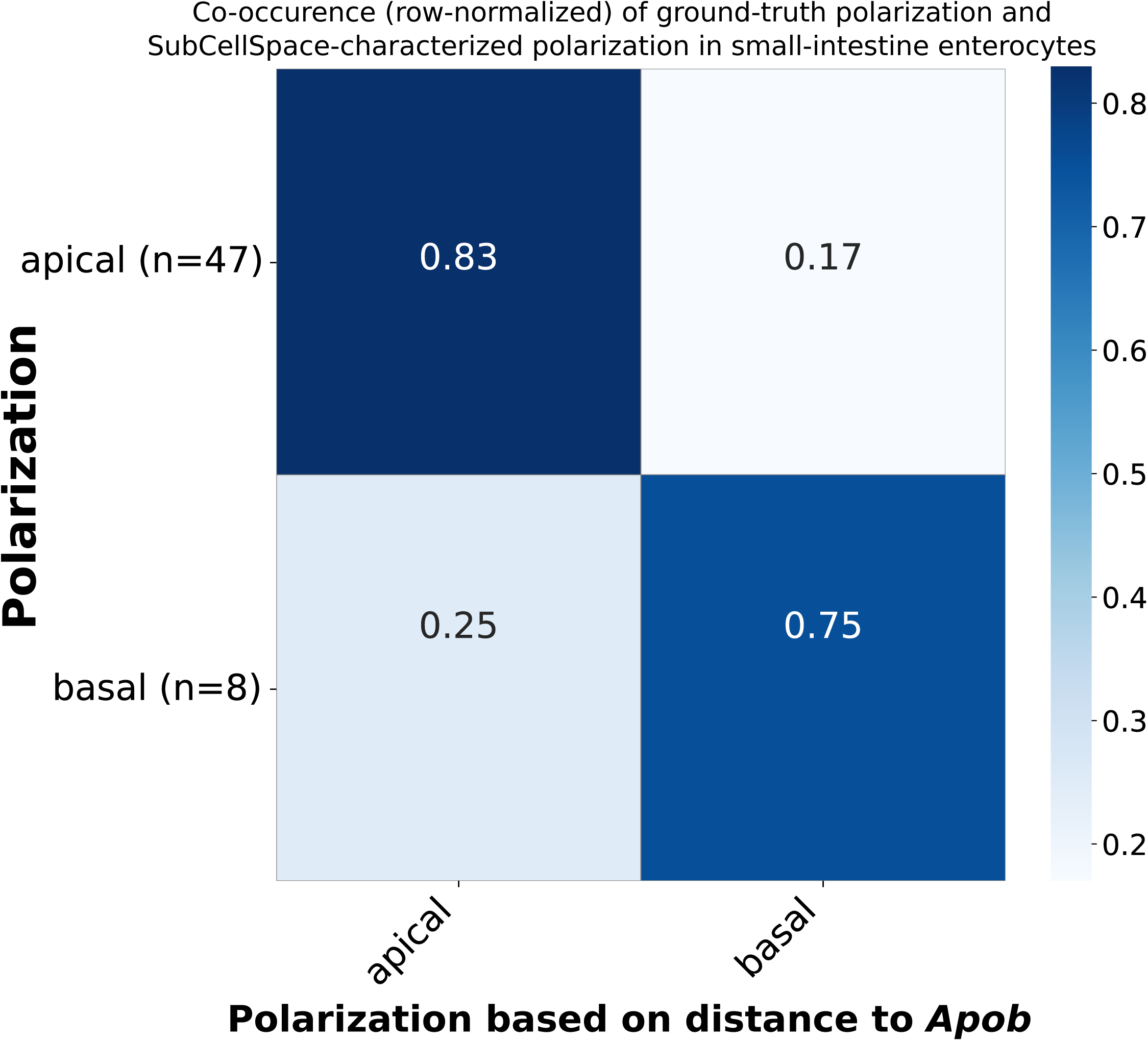
Apical/Basal classification of ground-truth polarized genes. Confusion matrix of flagging genes as apical/basal based on their cosine distance to apical marker Apob in SubCellSpace. Cell elements are row-wise normalized percentages. Total n is noted next to the true labels.

**Supp. Fig. 10:**
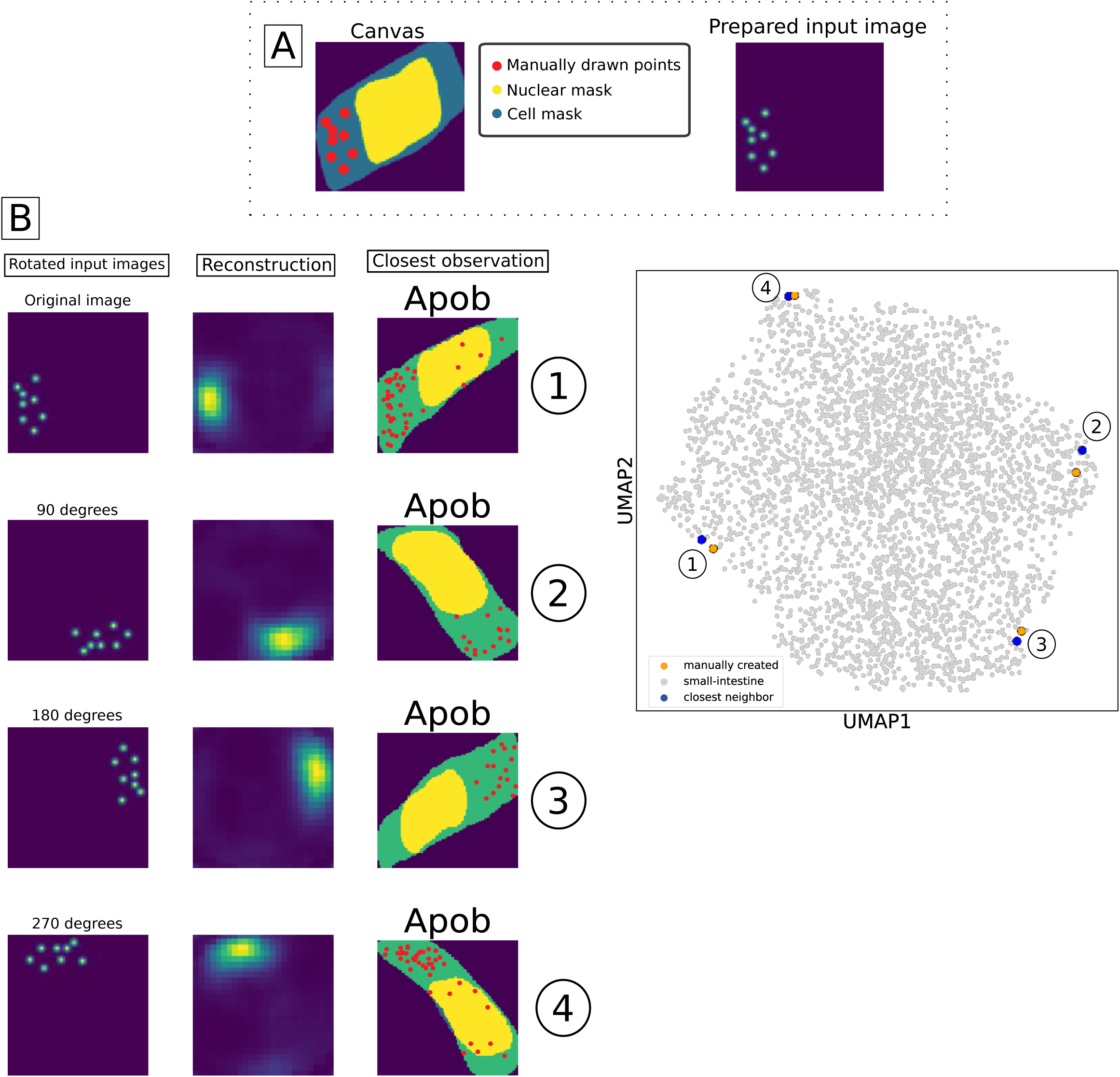
Retrieval of specific patterns without markers by manually drawing examples. This illustrates a workflow to retrieve examples of a specific pattern without access to a classification label or marker gene. **A.** Drawing of the input image. The canvas shows a random cell-mask (blue) + nuclear mask (yellow) from the available data. Dots (in red) are manually drawn in interactive Jupyter. The right image shows the prepared input image, which will be embedded into SubCellSpace. **B.** The prepared input image, along with 3 rotational augmentations, are embedded into subcellspace. The second column shows the reconstruction of these embeddings, and the third column shows the nearest real data neighbor within SubCellSpace. **C.** A UMAP of the SI MERFISH data with the 4 drawn input points co-embedded. The input points are indicated with orange dots and a number referring back to their order in B. Their closest real-data neighbor is indicated with a blue dot.

**Supp. Fig. 11:**
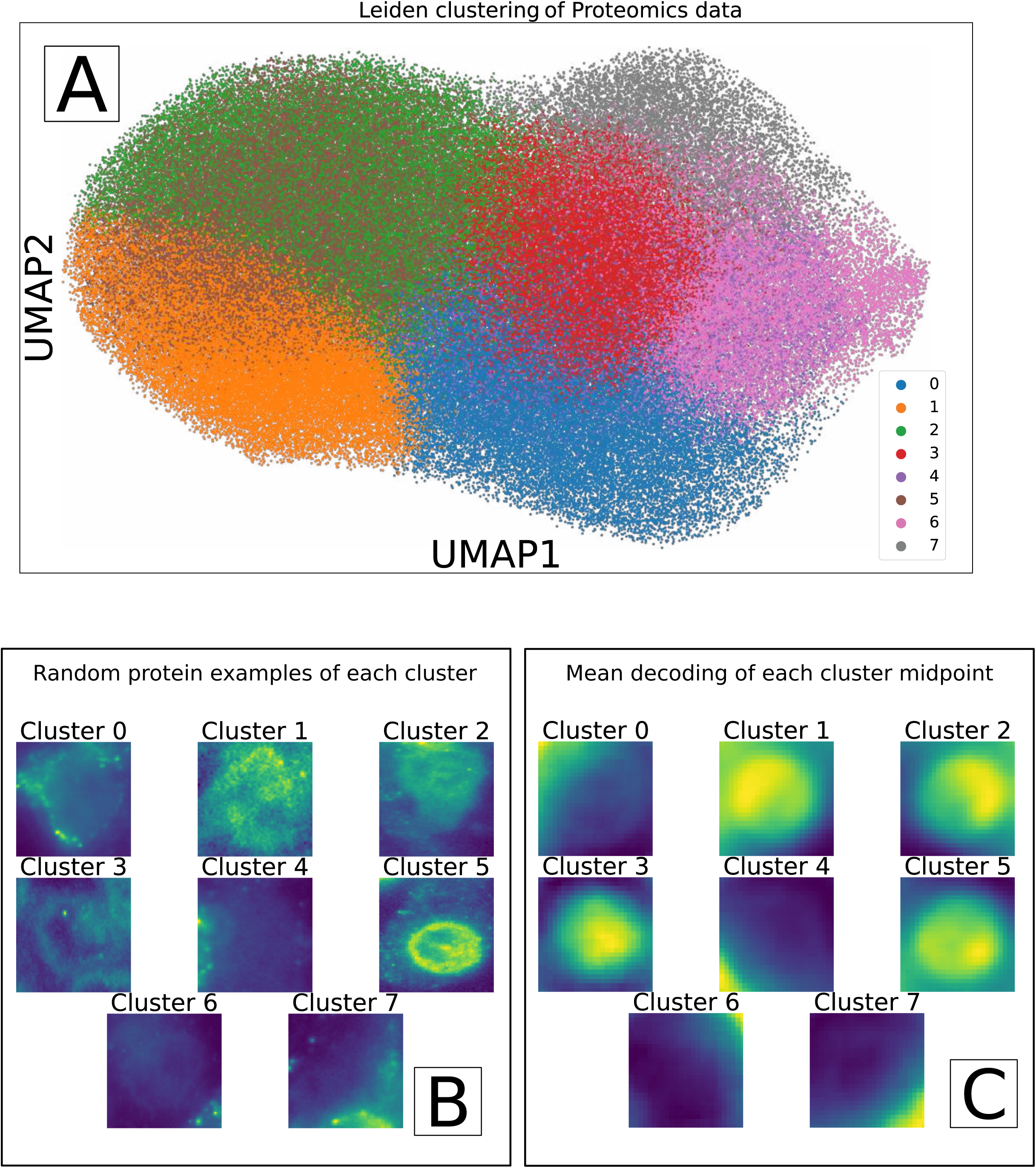
Transfer learned SubCellSpace of proteomics reveal clusters capturing heterogeneity in CD45R0 protein-staining. **A.** UMAP-representation the transfer-learned SubCellSpace of a CD45R0 protein-staining of a publicly available Vizgen co-expression MERFISH experiment. The UMAP is colored by a standard Leiden clustering. **B.** Representative example images of observations contained by each Leiden cluster. **C.** The reconstruction of each midpoint of all clusters.

## Notes

### Competing Interest Statement

The authors have declared no competing interest.

