## Supplementary Figures for "SubCellSpace: Automated characterization of subcellular mRNA localization patterns in spatial transcriptomics"

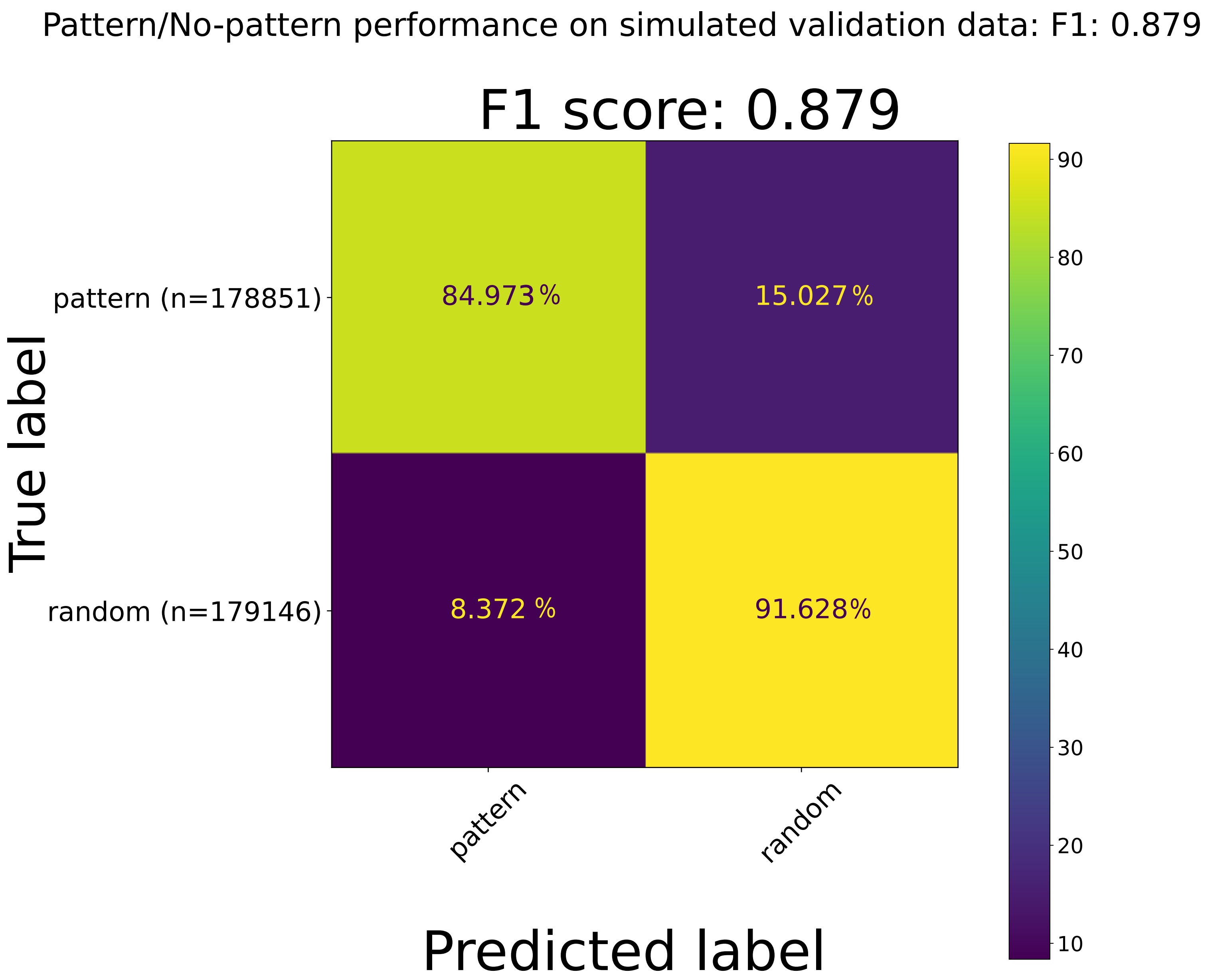


**Supp. Fig. 1: Pattern/no-pattern classification performance on simulated validation data.** Confusion matrix of the pattern/no-pattern classifier on the simulated validation data. Cell elements are row-wise normalized percentages. Total *n* is noted next to the true labels.

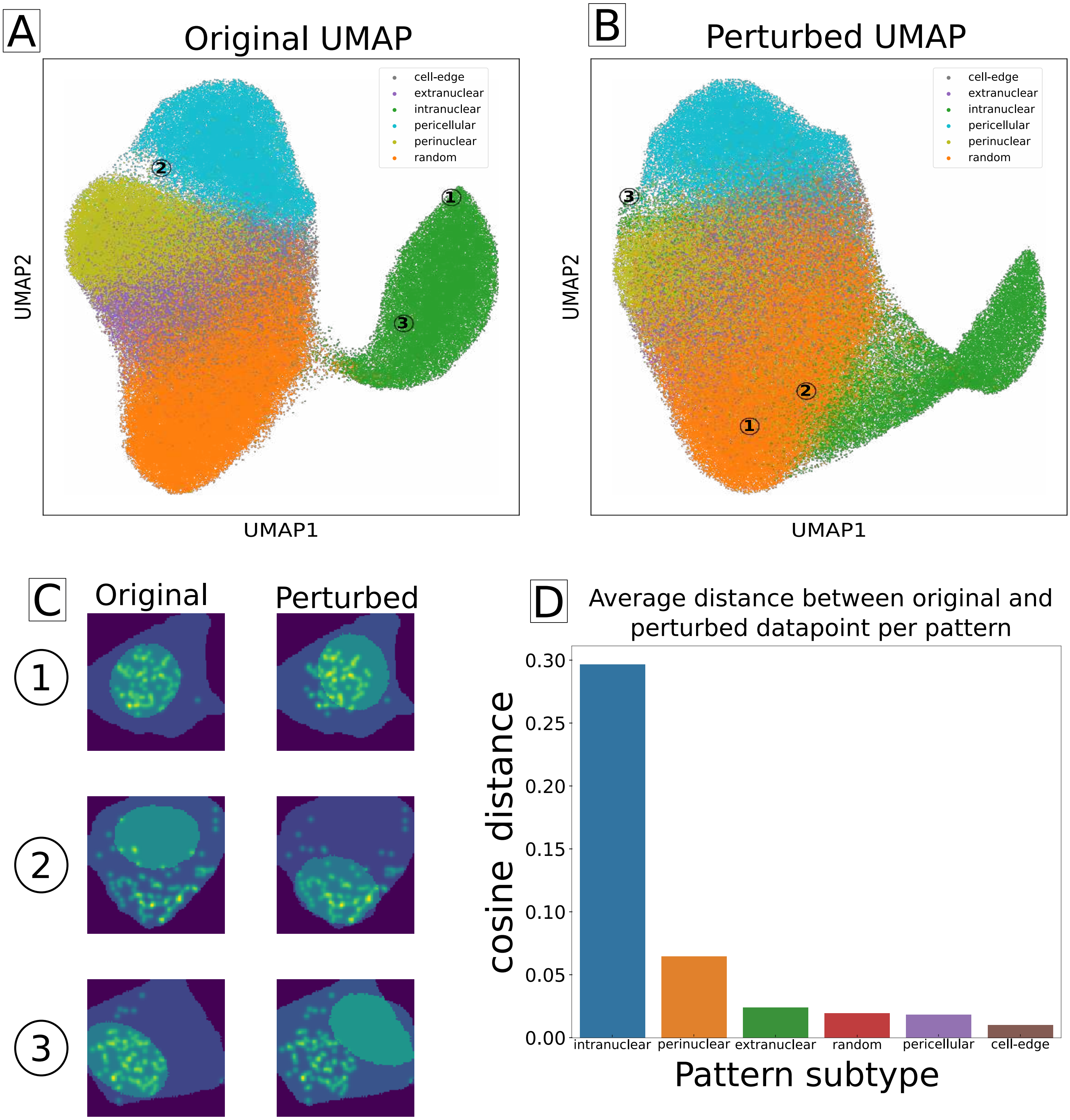


**Supp. Fig. 2: Perturbing the DAPI mask disrupts embeddings of nuclear-based patterns**
**A.** UMAP-representation of regular SubCellSpace, with 3 maximal perturbed observations indicated. **B.** UMAP-representation of a SubCellSpace after each DAPI-mask was randomly rotated. The same 3 observations are indicated with their new placement. **C.** 3 maximally perturbed observations, measured by pairwise cosine distance in multi-dimensional space. The left column is their original form, right is after perturbing the DAPI mask. **D**. Averaged distance per simulated pattern between original embeddings and perturbed embeddings.

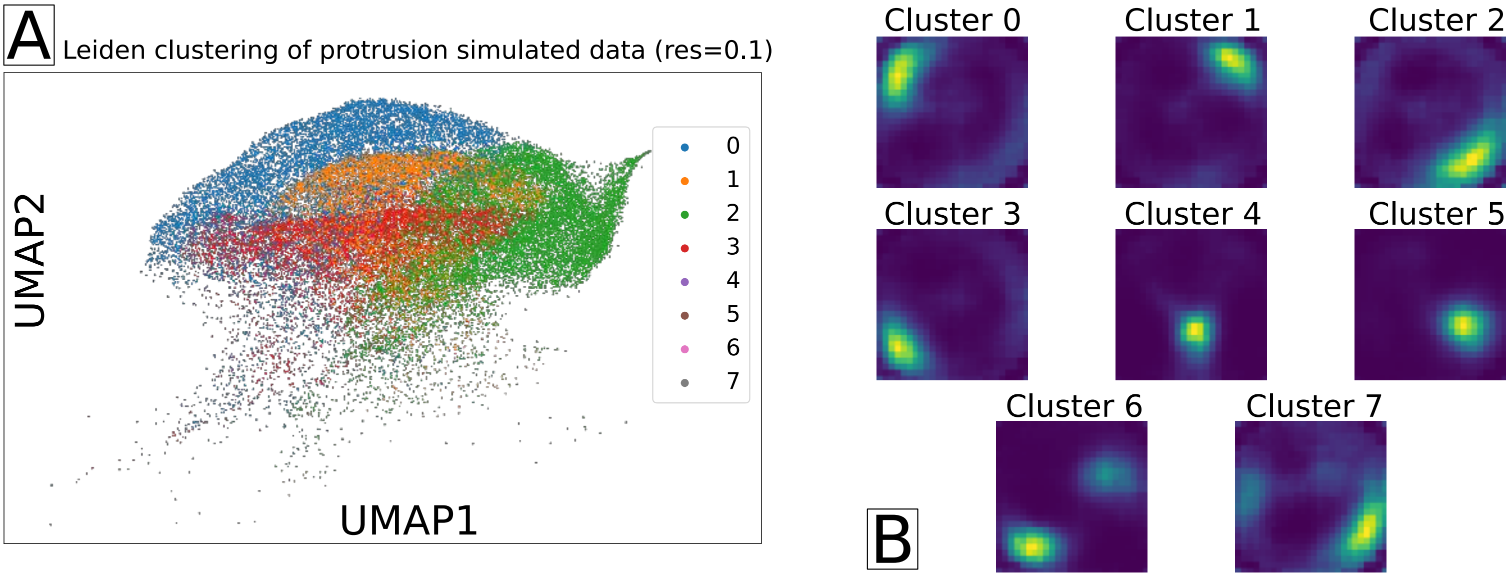


**Supp. Fig. 3: Subclustering of the unseen protrusion simulated pattern**
**A.** Cropped UMAP-representation of SubCellSpace showing only the unseen protrusion simulated observations, colored by cluster assignment by a Leiden clustering at resolution 0.1. **B.** Reconstruction of the midpoint of each leiden cluster.

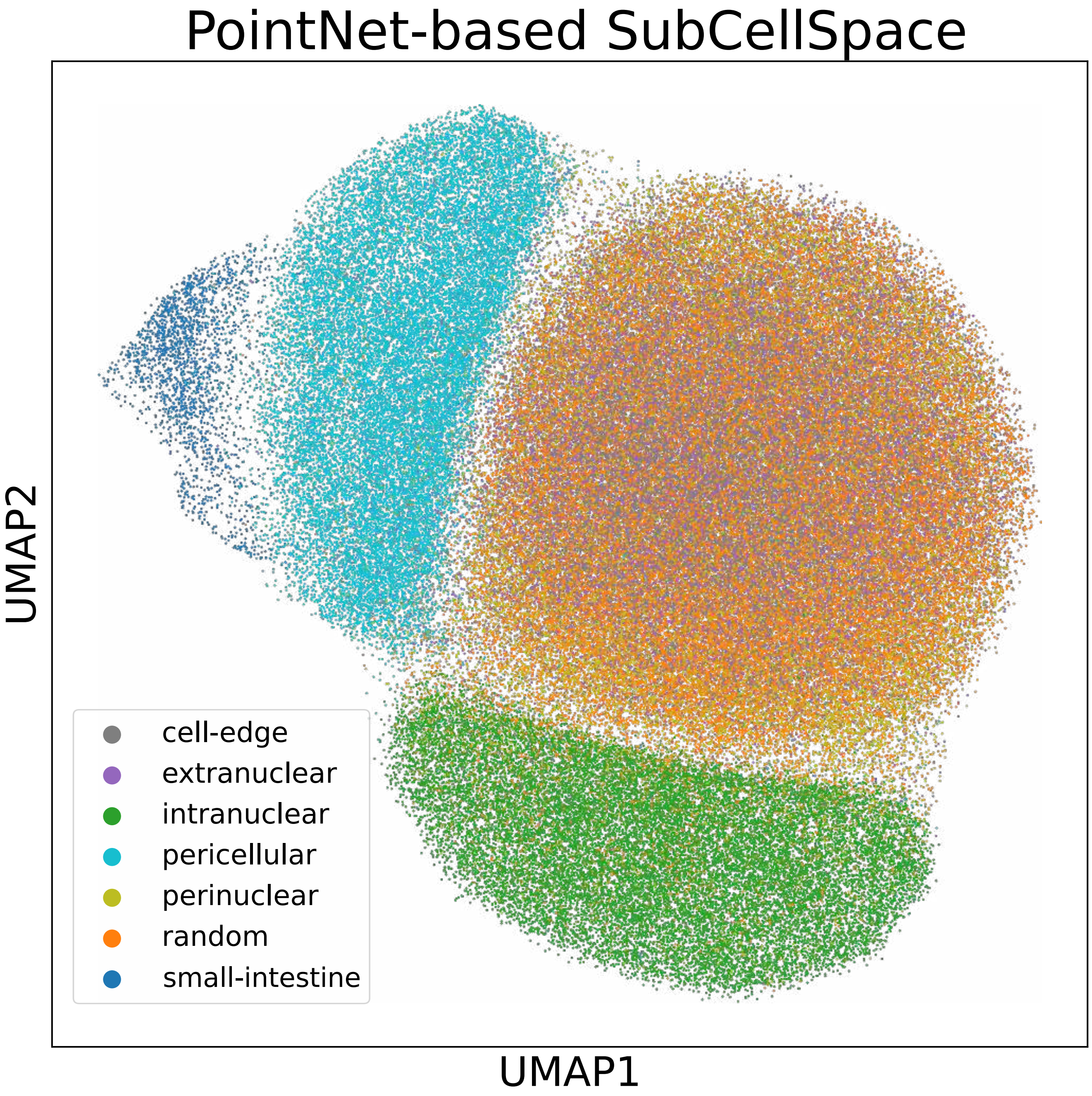


**Supp. Fig. 4: Point-cloud implementation of SubCellSpace**
UMAP-representation of the training data combined with the small-intestine MERFISH data. UMAP was calculated based on the point-cloud application-derived embeddings.

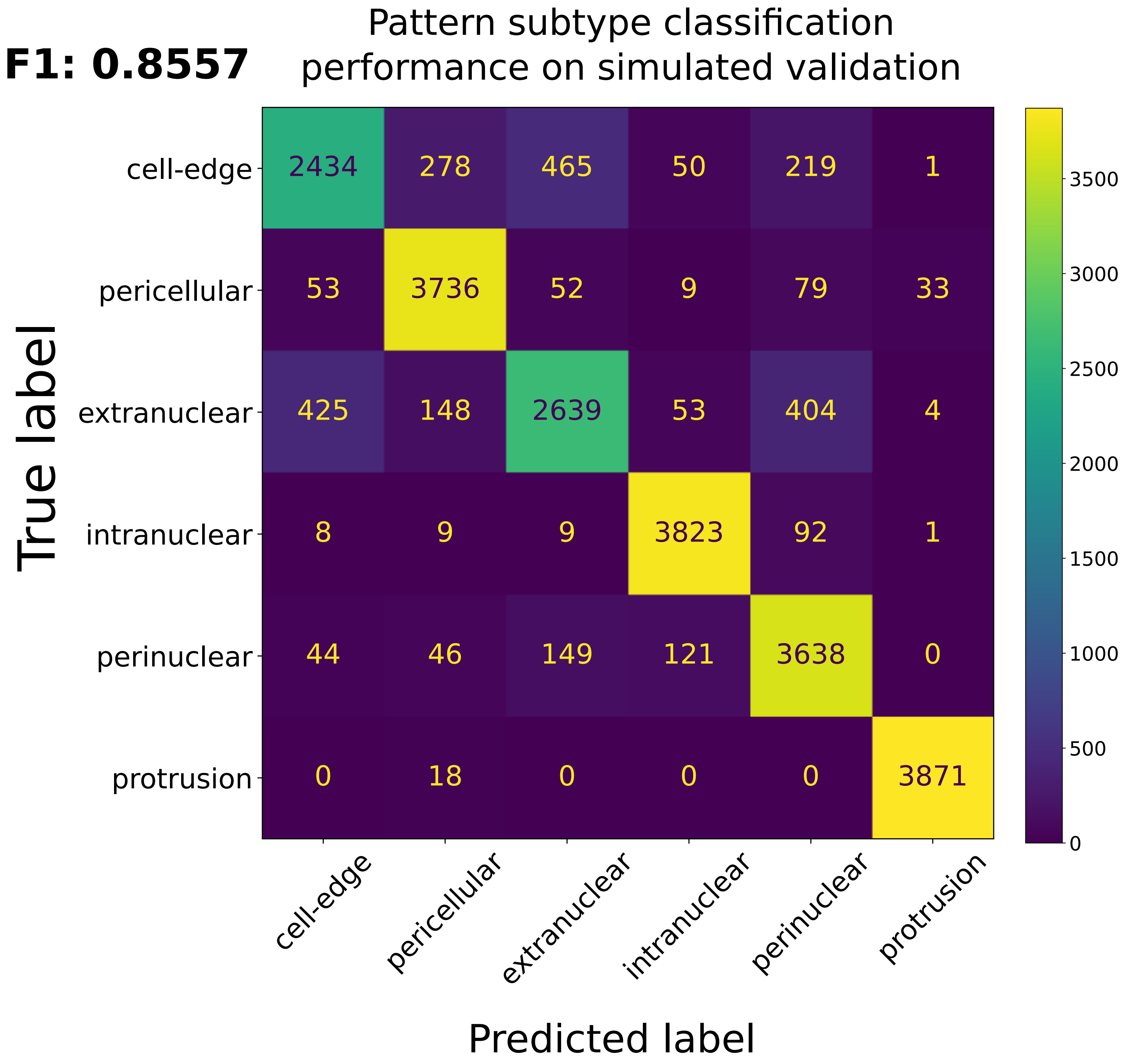


**Supp. Fig. 5: Pattern subtype classification performance on simulated validation data**
Confusion matrix of the pattern subtype classifier on the simulated validation data. Cell elements are absolute values.

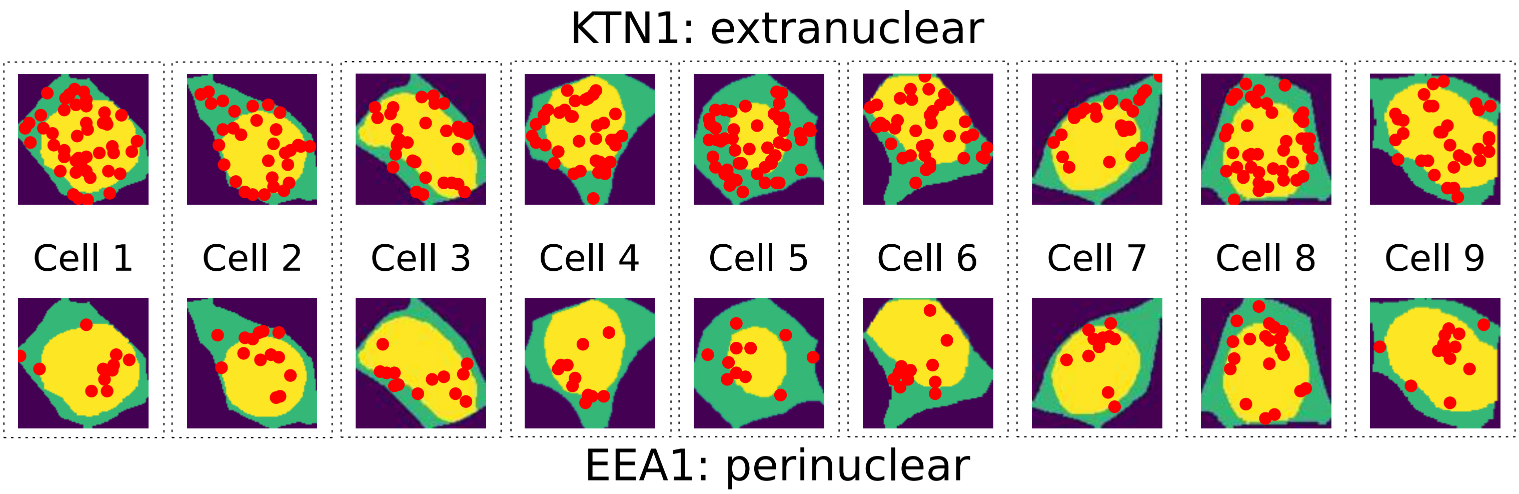


**Supp. Fig. 6: Heterogeneity in pattern-presentation of ERM-assigned genes.** Shown are 2 genes from the Xenium validation data, both of which have been assigned to be ERM-enriched by APEX-seq, but classified as different pattern subtypes by our workflow (KTN1 as extranuclear, and EEA1 as perinuclear). Columns show nine cells with maximal respective pattern-subtype scores and their mRNA localization in those cells.

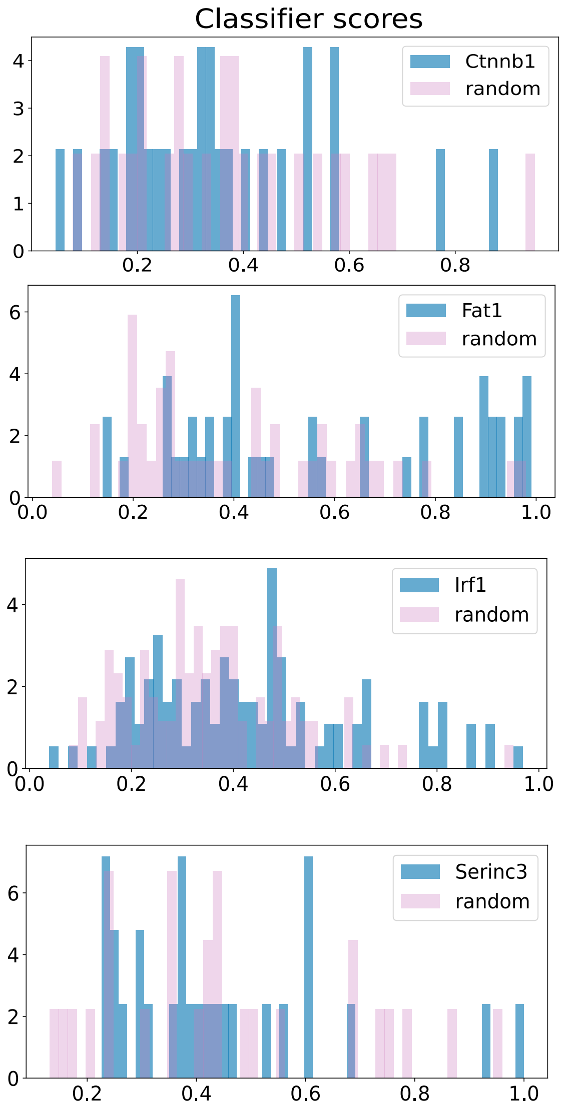


| | |
|---|---|
| gene_name | Ctnnb1 |
| n_obs | 28 |
| earth_dist | 0.042513 |
| pval | 0.773167 |
| bonferonni pval | 25.5145 |
| passes | False |
| passes_earth_thresh | False |

| | |
|---|---|
| gene_name | Fat1 |
| n_obs | 45 |
| earth_dist | 0.169145 |
| pval | 0.025096 |
| bonferonni pval | 0.828175 |
| passes | False |
| passes_earth_thresh | True |

| | |
|---|---|
| gene_name | Irf1 |
| n_obs | 99 |
| earth_dist | 0.07986 |
| pval | 0.014823 |
| bonferonni pval | 0.489166 |
| passes | False |
| passes_earth_thresh | True |

| | |
|---|---|
| gene_name | Serinc3 |
| n_obs | 27 |
| earth_dist | 0.052226 |
| pval | 0.935662 |
| bonferonni pval | 30.876854 |
| passes | False |
| passes_earth_thresh | False |

**Supp. Fig. 7: Pattern-probability distributions of False Negatives of the small-intestine MERFISH data.** Pattern-probability distributions for each gene of the small-intestine dataset that was falsely classified as not-pattern-presenting, together with its randomized distribution. Noted for each gene is their EMD, p-val, adjusted p-val, and whether those values pass their respective thresholds.

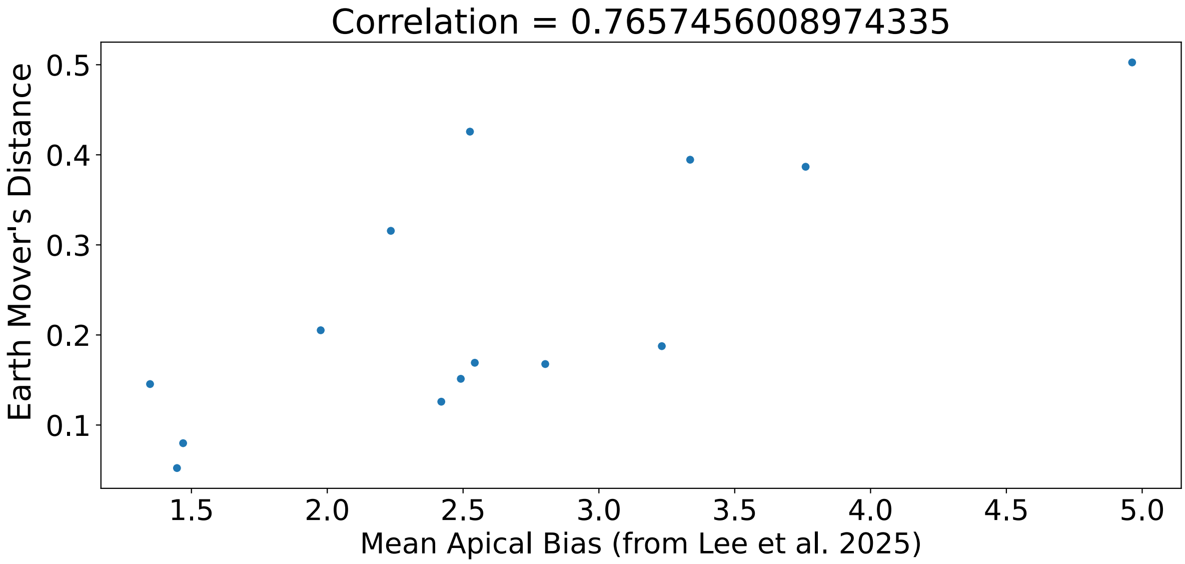


**Supp. Fig. 8: EMD's of apical genes correlate to their apical bias**
A scatterplot showing the correlation for each ground-truth apically polarized gene from the SI MERFISH data. Each dot is a gene, where the x-axis represents its noted apical bias as reported in Lee et al. 2025. The y-axis is this gene's EMD compared to its randomized counterpart. Correlation was calculated as the Pearson's correlation between the two metrics.

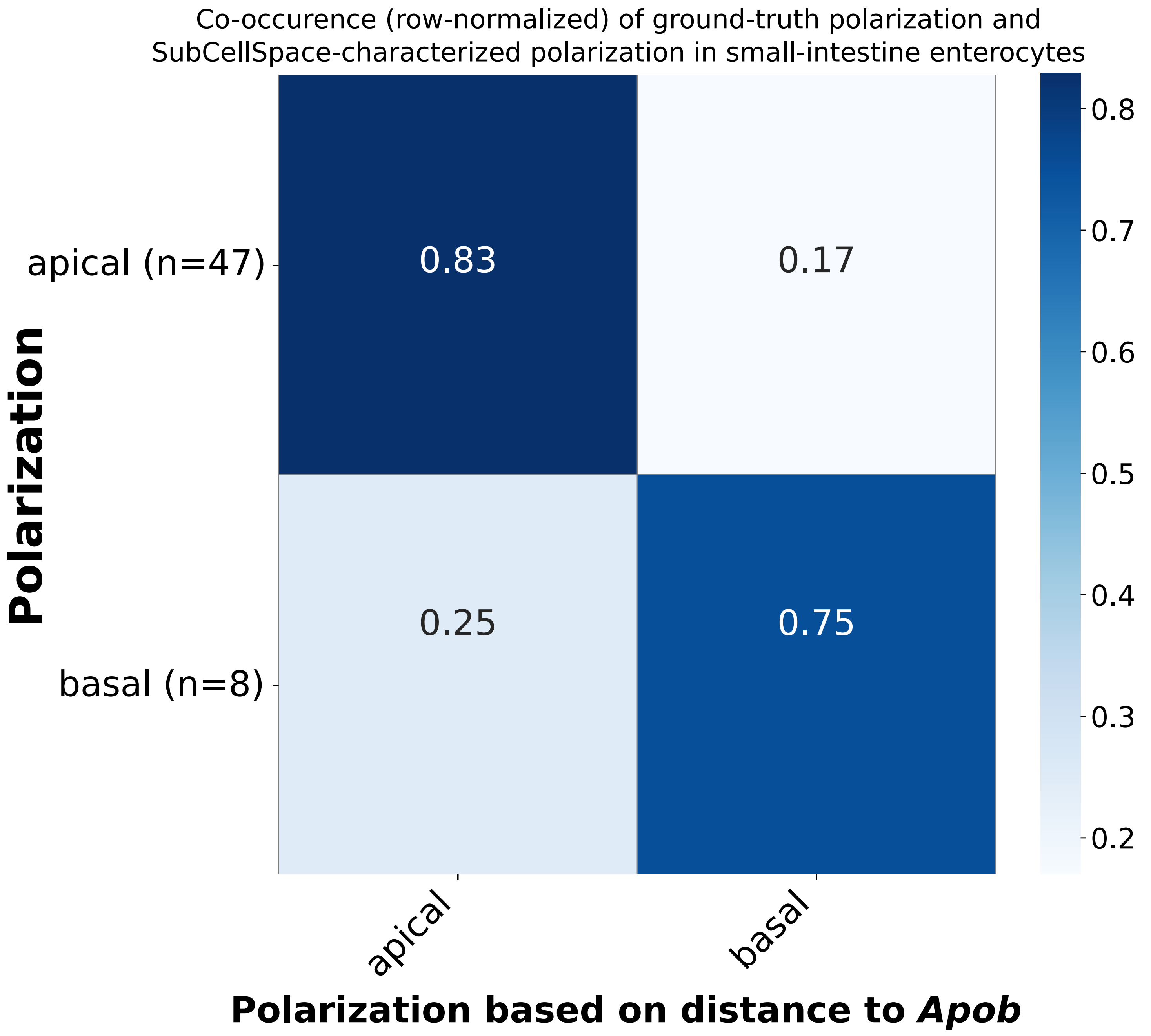


**Supp. Fig. 9: Apical/Basal classification of ground-truth polarized genes**
Confusion matrix of flagging genes as apical/basal based on their cosine distance to apical marker Apob in SubCellSpace. Cell elements are row-wise normalized percentages. Total n is noted next to the true labels.

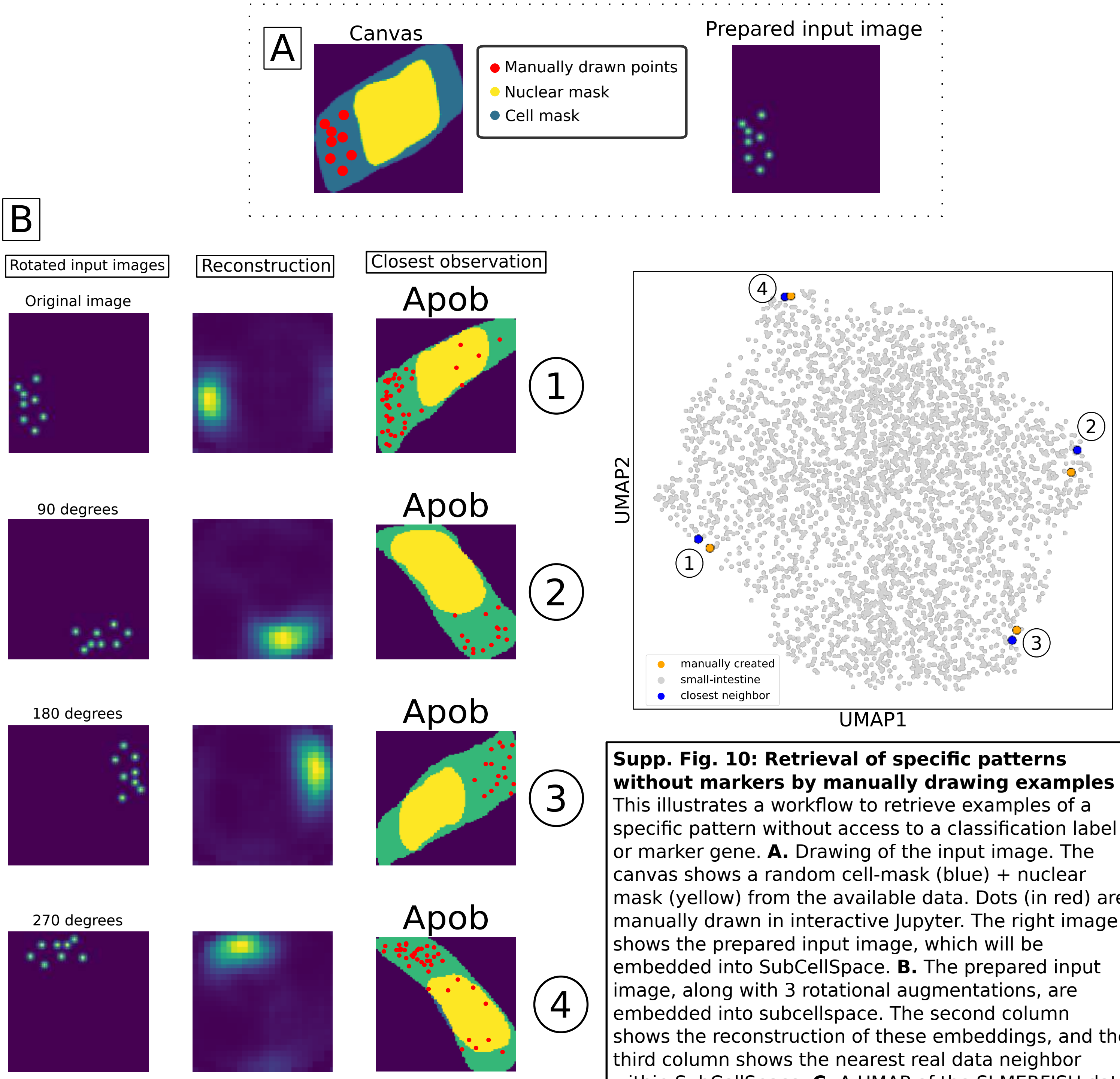


**Supp. Fig. 10: Retrieval of specific patterns without markers by manually drawing examples** This illustrates a workflow to retrieve examples of a specific pattern without access to a classification label or marker gene. **A.** Drawing of the input image. The canvas shows a random cell-mask (blue) + nuclear mask (yellow) from the available data. Dots (in red) are manually drawn in interactive Jupyter. The right image shows the prepared input image, which will be embedded into SubCellSpace. **B.** The prepared input image, along with 3 rotational augmentations, are embedded into subcellspace. The second column shows the reconstruction of these embeddings, and the third column shows the nearest real data neighbor within SubCellSpace. **C.** A UMAP of the SI MERFISH data with the 4 drawn input points co-embedded. The input points are indicated with orange dots and a number referring back to their order in B. Their closest real-data neighbor is indicated with a blue dot.

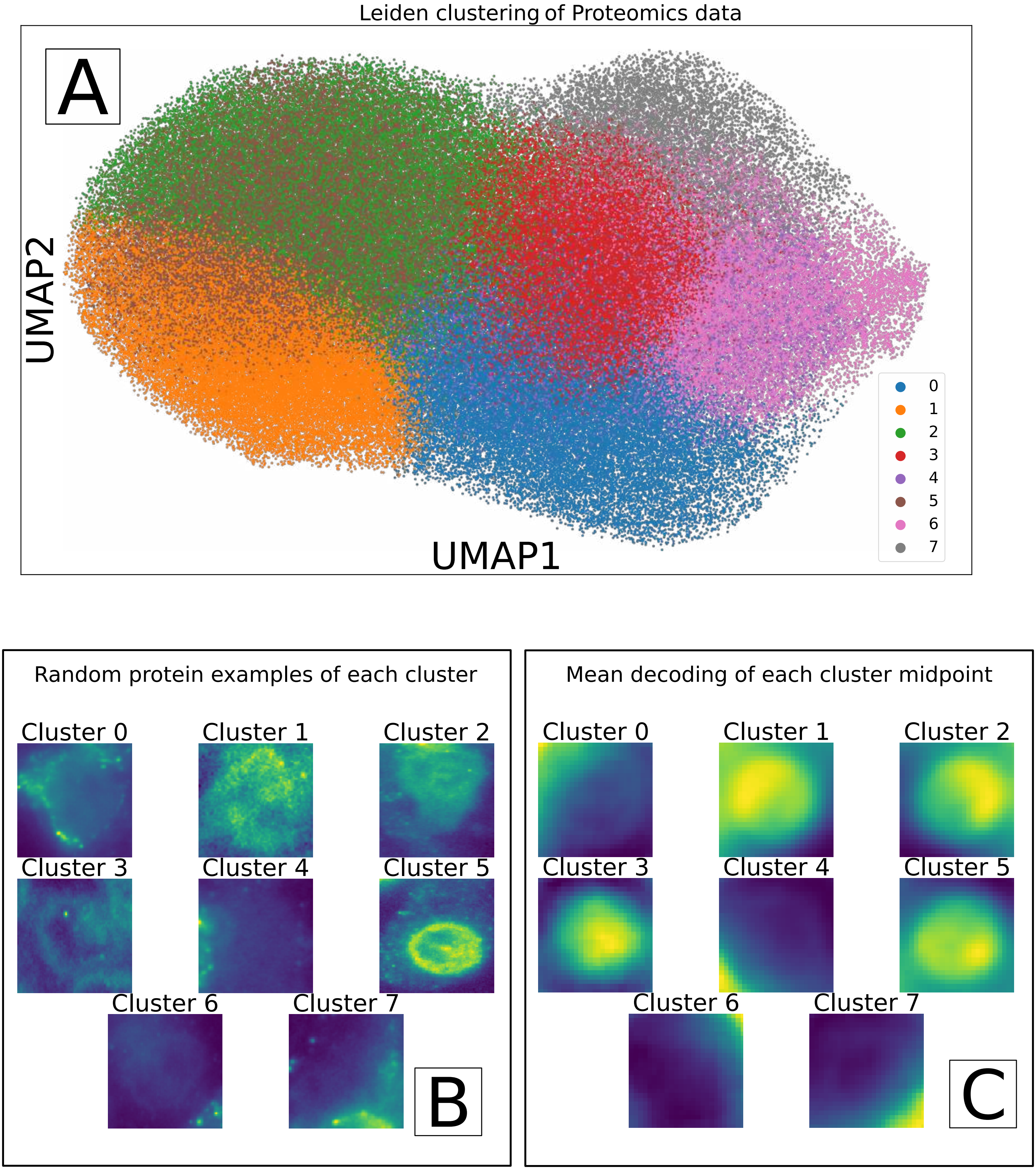


**Supp. Fig. 11: Transfer learned SubCellSpace of proteomics reveal clusters capturing heterogeneity in CD45R0 protein-staining**
**A.** UMAP-representation the transfer-learned SubCellSpace of a CD45R0 protein-staining of a publicly available Vizgen co-expression MERFISH experiment. The UMAP is colored by a standard Leiden clustering. **B.** Representative example images of observations contained by each Leiden cluster. **C.** The reconstruction of each midpoint of all clusters.
